# Virtual experiments bridge sequence and microscopy with generative models

**DOI:** 10.64898/2026.09.13.751243

**Authors:** Dihan Zheng, Kibeom Hong, Bo Huang

**Affiliations:** Department of Pharmaceutical Chemistry, University of California, San Francisco, San Francisco, CA 94143; Department of Biochemistry and Biophysics, University of California, San Francisco, San Francisco, CA 94143; Biohub San Francisco, San Francisco, CA 94158

## Abstract

Large-scale screening and mapping efforts have produced vast libraries of perturbation-readout data. Converting these measurements into mechanistic insights requires models that link perturbation and genetic input to phenotypes, i.e., labels from experimental readouts, which are usually task specific. We propose a different, virtual experiment modeling approach: train generative models to recreate readouts conditioned on the experimental context, and then let established downstream models extract phenotypes from the synthetic data. As an illustrative case, we develop a bidirectional sequence-image generative framework, CELL-FM, that maps protein sequence and cellular context to fluorescence microscopy images and back, enabling *in silico* localization prediction, image-conditioned functional motif analysis and generation, and large-scale virtual mutagenesis revealing the amino acid features controlling condensate formation of intrinsically disordered peptides. This approach decouples representation learning from task-specific annotation, reuses rich experimental modalities across many downstream tasks, and preserves the spatial and organizational detail that hand-crafted labels often discard.

## Introduction

High-throughput screening and mapping technologies now generate vast amounts of sequence- or perturbation-linked measurements. Typical predictive modeling pipelines compress those measurements into task specific phenotype labels before learning, such as the depiction of gene functions using Gene Ontology (GO) terms^1–3^, the annotation of cell type and cell state from single-cell sequencing readout^4–6^, and the description of protein localization based on predefined subcellular compartments^7–9^. Building separate labeled datasets for every biological question, however, is inefficient, making it harder to transfer models across questions. Moreover, labels can be lossy: many phenotypes are continuous, context-dependent, and spatially structured, so reducing them to discrete categories discards information. Here, instead of training end-to-end predictors for curated labels, we showcase training generative models that learn to reproduce the original experimental readouts of the assay. Once a model can synthesize realistic experimental outputs conditioned on genotype, perturbation, context, etc., downstream phenotyping becomes a separate step that utilizes existing analysis pipelines, classifiers, or feature extractors established for the real assay. This two-stage approach has three practical advantages: (1) it leverages the full richness of the experimental readouts rather than forcing early compression into labels; (2) it enables reuse of a single generative model for many downstream tasks without retraining; and (3) it supports controlled *in silico* experiments that recapture biological variabilities while factoring out other, unnecessary variabilities present in real screens.

We illustrate this paradigm with the modeling between protein sequence inputs and cellular fluorescence microscopy readouts, of which the experiments have now become highly scalable with large-scale antibody generation^10,11^, genome engineering^12^, ORFome libraries^13,14^, and optical pooled screening (OPS) technologies^15,16^. Fluorescence images encode subcellular targeting, mesoscale organization, and cell-to-cell variability: features that are difficult to capture with simple labels but are directly usable by image-based analysis tools^17–19^. By training a conditional generative model to map protein sequence and cellular context to realistic fluorescence images, we create a virtual microscopy platform, CELL-FM, that supports sequence-conditioned image prediction as well as image-conditioned sequence sampling. Because downstream phenotype quantification is performed by the same image analysis pipeline used on experimental data, the virtual experiments supported by CELL-FM remain directly comparable to wet-lab assays and can be used to prioritize hypotheses and guide follow-up experiments.

CELL-FM is a unified generative framework that jointly models fluorescence microscopy images as a continuous distribution and protein sequences as a discrete distribution (Fig. 1a). Rather than learning a single deterministic mapping between sequence and image as in several existing sequence-to-image models^20–22^, CELL-FM adopts a generative formulation that we demonstrated previously^23^ to capture the intrinsic stochasticity of biological images and infer from underdetermined contextual input. Trained with public large-scale microscopy mapping data including the Human Protein Atlas (HPA)^10,11^, OpenCell^12^, and CondenSeq^24^, CELL-FM can reproduce realistic single-cell variability and localization patterns, reveal sequence determinants through large-scale *in silico* mutations, and generate functional localization motifs when constrained by image phenotypes. More generally, this work shows how modeling experimental readouts rather than pre-defined labels creates a flexible, reusable foundation for biological discovery, exemplified by our characterization of amino acid determinants of unexpected biomolecular condensate formation phenotypes.

**Figure 1:**
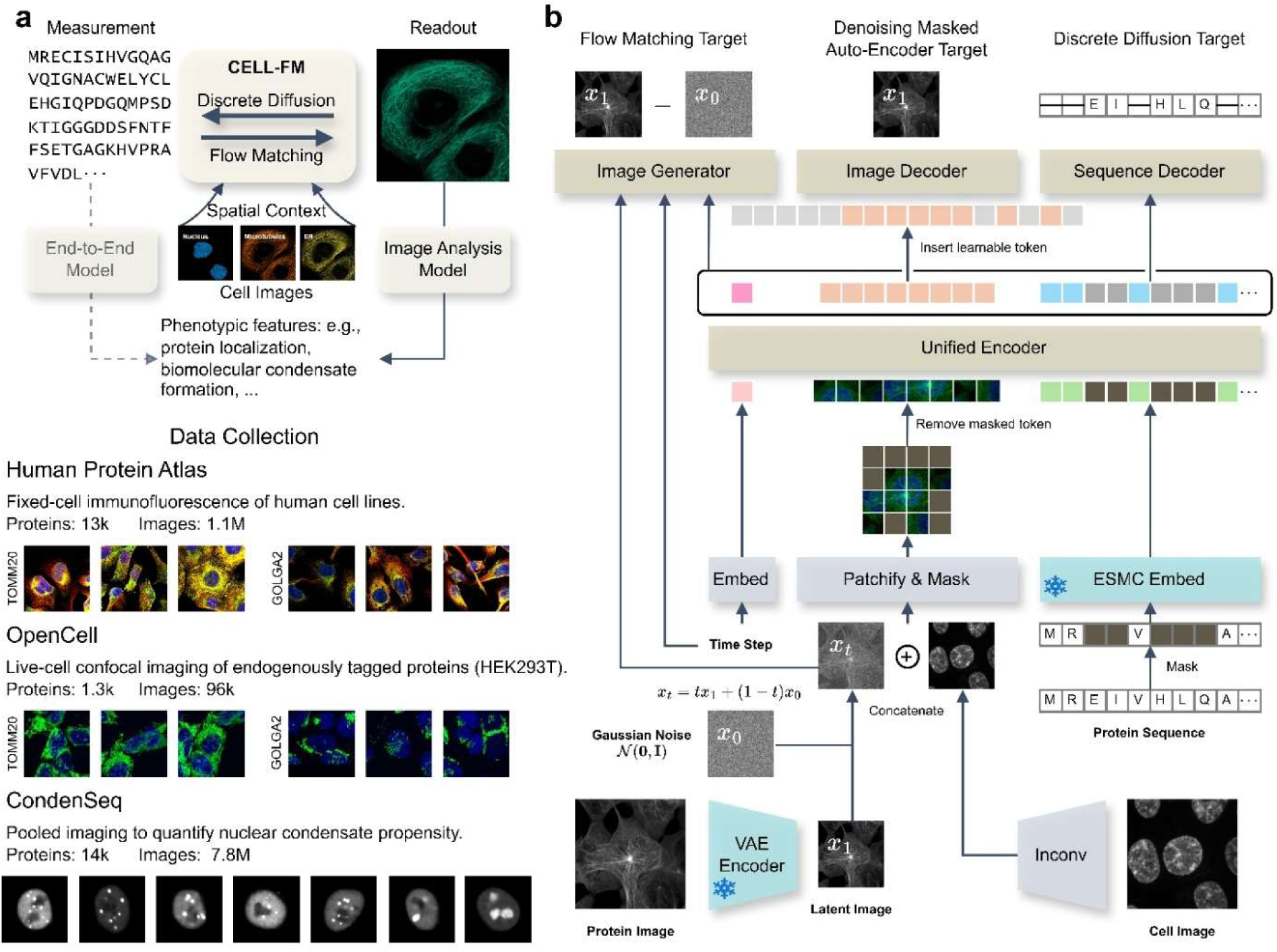
Data resources and model architecture of CELL-FM for sequence–image generation. **a.** The task overview: CELL-FM is a unified generative framework that models protein images (continuous diffusion) and protein sequences (discrete diffusion), enabling conditional, bidirectional generation. Given cell images as conditional inputs, the model can generate protein fluorescence images from protein sequences and, conversely, propose protein sequences consistent with observed fluorescence patterns. In contrast to conventional end-to-end predictors that map sequence directly to a predefined functional label, CELL-FM produces an interpretable intermediate imaging modality that supports a broad range of downstream analyses. **b.** Model and training objectives.

## Results

### Bidirectional generative model bridges protein sequence and cellular imaging

CELL-FM is a generative framework that jointly models fluorescence microscopy images as a continuous distribution and protein sequences as a discrete distribution (Fig. 1a). We adopt a unified encoding and separate decoding architecture (Fig. 1b). All inputs are first mapped into a continuous token space: multichannel images are patchified into image tokens, and protein sequences are converted into sequence tokens from a pretrained protein language model embedding. These tokens are then integrated by a unified encoder, where cross-modal information is fused into a shared representation. More specifically, it ingests patch tokens from the target protein fluorescence channel with accompanying cell context channels and sequence tokens from ESMC-600M^25^. Cell context is input as a conditioning signal by concatenating context channels with the protein channel before patchification and masking, enabling the encoder to learn how sequence-driven localization cues interact with morphology dependent spatial constraints.

The output heads act as modality-specific interpreters that read from the shared representation to produce different outputs under different training objectives. Three decoders branch from this shared backbone, each aligned to a distinct generative role. For protein image generation, we operate in the latent space of a variational autoencoder^26,27^ (VAE) encoder and train an image generator with a flow matching^28^ objective on interpolated latent *x*_t_ = *tx*_1_ + (1 − *t*)*x*_0_, where *x*_0_ ∼ *N*(0, **I**) and *x*_1_ is the target latent. In parallel, an image decoder is trained with a denoising masked autoencoder^29,30^ objective to reconstruct masked image tokens, encouraging robust and transferable representations. Finally, for sequence generation, a sequence decoder is trained with a discrete diffusion^31,32^ objective over protein tokens, allowing the model to propose amino acid sequences conditioned on the same unified representation derived from images and cell context. This unified generative design enables both sequence-to-image generation and image-to-sequence design within a single model family, providing a foundation for downstream virtual microscopy applications.

### CELL-FM generates high fidelity images for unseen proteins

The first capability of CELL-FM is performing virtual microscopy experiments: given a protein sequence and a cellular context image, the model synthesizes plausible cellular fluorescence images that can be subsequently used to characterize how the protein might localize inside a cell (Fig. 2a). To learn how sequence-encoded localization cues interact with cellular morphology, we trained CELL-FM with the HPA subcellular resource^10,11^, a large, fixed human cell line immunofluorescence dataset providing both protein fluorescence channels and three contextual markers (nucleus, microtubules, ER). We curated 12,894 proteins with matched sequences, each associated with a few hundred single-cell crops, yielding 1.1 million single-cell images in total. We hold out 1,200 proteins as an unseen test set and train on the remaining proteins. Training proceeds in two stages. In the representation pretraining stage, we jointly optimize all three objectives using an image masking ratio of 0.5, encouraging the unified encoder to learn transferable multimodal representations. In the subsequent generative finetuning stage, we optimize the flow matching and discrete diffusion objectives with the image masking ratio set to 0, focusing capacity on high fidelity generation of fluorescence images.

**Figure 2:**
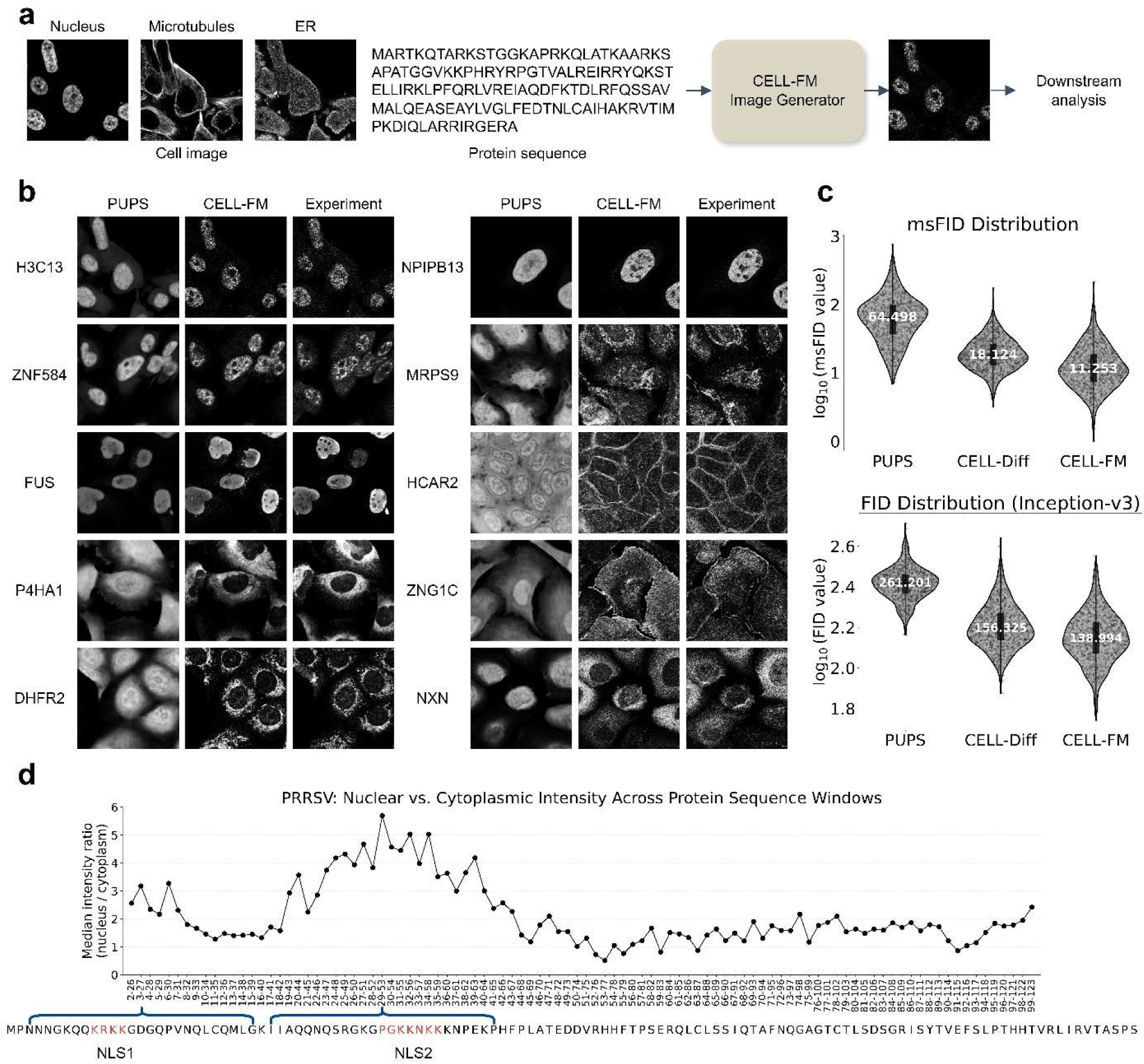
CELL-FM predicts protein localization from sequence and cell context. **a.** Overview of the CELL-FM image-prediction pipeline. **b.** Comparing fluorescence images predicted by CELL-FM and PUPS with experimental images for proteins in the test set. **c.** Quantitative evaluation of image-generation fidelity for the test set. **d.** NLS identification by CELL-FM. A sliding window of 25 amino acids is applied across the PRRSV protein sequence. For each peptide window, CELL-FM generates 64 images and computes a nuclear enrichment score as the ratio of mean fluorescence density inside versus outside the nucleus. The median score across the 64 generated images is used as a robust estimate of each peptide’s nuclear-transport potential.

HPA-trained CELL-FM learns to generate high fidelity fluorescence images for unseen proteins. When provided with a test set sequence and a reference cell morphology, the model produces a distribution of single-cell images that capture the stereotyped localization pattern of the protein. Qualitatively, the generated images recover fine spatial features such as nucleolar enrichment, filamentous cytoskeletal association, and organelle specific targeting, while avoiding the over-smoothed, mean-like predictions typical of deterministic models such as PUPS^22^ (Fig. 2b). Quantifying the fidelity of the generated images, however, is challenging for common pixel-wise image similarity metrics such as Pearson correlation coefficient (PCC) and peak signal-to-noise ratio (PSNR), which actually reward mean-like predictions and global intensity alignment, while remaining insensitive to higher-order semantic attributes that define protein localization (Supplementary Fig. 2a). Therefore, we adopt distributional similarity, including Fréchet Inception Distance^33^ (FID) and Kernel Inception Distance^34^ (KID), which operate by embedding images into a feature space and comparing the resulting feature distributions. Besides standard FID relying on embeddings from natural images such as Inception-v3^35^, we also trained a microscopy-specific embedding model based on a vision transformer^36^ (ViT) to establish the cellular microscopy adapted metrics msFID/msKID (Supplementary Fig. 2b). All distributional metrics show a substantial improvement of CELL-FM over PUPS and also over previous generative model CELL-Diff^23^ (Fig. 2c). These results indicate that CELL-FM better matches the single-cell variability and fine scale spatial structure of protein localization for unseen proteins.

CELL-FM generated images can then be fed to standard image analysis pipelines to measure protein localization. We demonstrate this application by identifying nuclear localization signals (NLSs) from the nucleocapsid protein of porcine reproductive and respiratory syndrome virus (PRRSV). By sliding a 25 a.a. window across the protein and generating 64 images for each peptide segment, we computed median nuclear-enrichment scores directly from the synthetic images (Fig. 2d). The resulting profile correctly highlights the two known NLS regions^37^, showing that CELL-FM can reveal functional motifs by converting sequence hypotheses into virtual fluorescence readouts. This illustrates how virtual microscopy experiments can support sequence-level screening without requiring explicit phenotype labels.

### CELL-FM generates functional NLS/NES motifs

As a model linking the amino acid sequence of proteins and cellular fluorescence images, CELL-FM is also capable of generatively sampling sequence motifs that lead to a specific subcellular localization pattern or cellular phenotype as described by an input fluorescence image. In this way, the consensus features of motifs can be statistically determined from the generated population, and *de novo* functional motif sequences can also be created. For this application, we fine-tuned the HPA-pretrained model using the full HPA dataset to leverage all available data.

To demonstrate this application, we focused on nuclear localization signals (NLSs) and nuclear export signals (NESs), which are well represented in the HPA training data. Here, instead of using categorical labels of localization phenotypes or specifically mining the human proteome for NLS/NES, we simply used a nuclear-enriched fluorescence image and a nuclear-depleted image plus their corresponding contextual images (Fig. 3a), asking the model to generate candidate sequences with lengths ranging from 10 to 25 amino acids. For each length, we sampled 20 sequences and performed amino acid frequency analysis. The generated NLS and NES pools showed clear compositional differences (Fig. 3b). NLS candidates were enriched for basic residues, including lysine (K) and arginine (R), whereas NES candidates were enriched for hydrophobic residues, including valine (V), leucine (L), and isoleucine (I). Relative to the proteome baseline, generated NLS motifs showed strong enrichment of positively charged residues, particularly K and R, together with depletion of several neutral or hydrophobic residues (Fig. 3b). In contrast, generated NES motifs were enriched for hydrophobic residues, including V, L, I, and W, and depleted for residues less associated with NES activity (Fig. 3b). These sequence-level trends are consistent with canonical localization-signal chemistry^38–40^.

**Figure 3:**
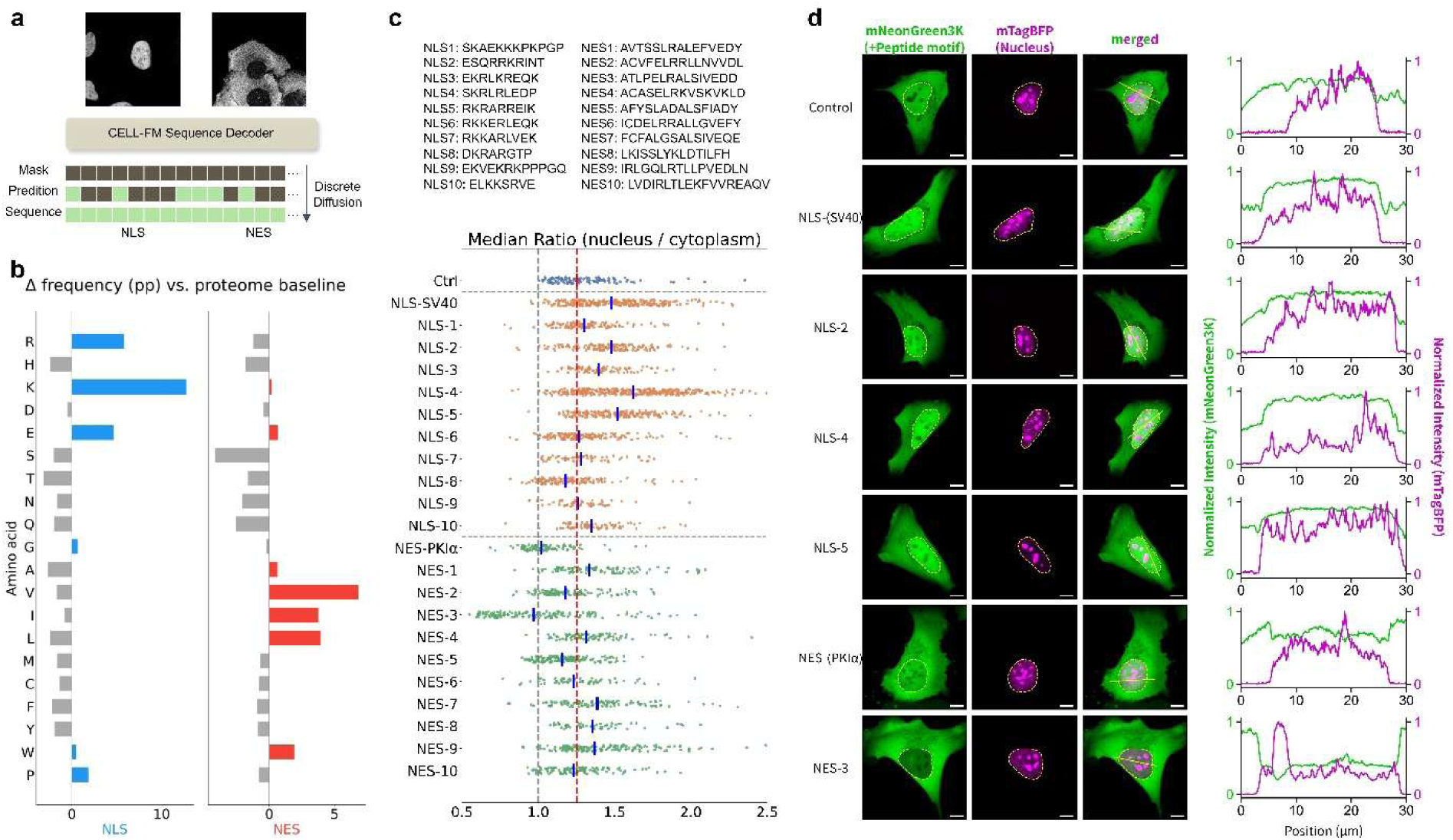
CELL-FM generates functional nuclear localization and export signals. **a.** Schematic of the sequence-design workflow. Given an input cellular fluorescence image, CELL-FM sequence decoder generates candidate localization motifs through discrete diffusion. **b.** Per-residue frequency differences of generated NLS and NES sequences relative to the proteome baseline. Positive values indicate residues enriched in the generated motif. **c.** The sequence of the 10 candidate NLS and NES sequences, and quantitative screening of generated motifs using the nuclear-to-cytoplasmic fluorescence intensity ratio. Several generated NLSs, including NLS-2, NLS-4, and NLS-5, promote nuclear enrichment comparable to the canonical SV40 NLS, while NES-3 drives nuclear exclusion comparable to the PKIα NES. Each point represents a single cell; vertical reference lines indicate control or benchmark motif performance. **d.** Representative fluorescence images and line-intensity profiles for selected motifs.

To create new NLS/NES, we added a screening step after sequence generation, using AlphaFold3^41^ to model the interaction between the generated sequences (5 to 25 a.a. and 40 sequences for each length) with the corresponding transport receptors: importin KPNA1 for NLS candidates and exportin CRM1 for NES candidates. We used the ipTM score^42^ to prioritize candidates with plausible receptor binding. We selected 10 NLS motifs and 10 NES motifs for experimental validation by expressing them in human U2OS cells as N-terminal fusions to the fluorescent protein mNeonGreen3K. Using a co-expressed 3xNLS-mTagBFP nuclear marker for image segmentation, we assessed the nuclear-to-cytoplasmic ratio of mNeonGreen fluorescence intensity (Fig. 3c). The 3 top-performing generated NLSs (NLS-2, NLS-4 and NLS-5) give nuclear enrichment of mNeonGreen3K comparable to the widely used SV40 NLS, with higher nucleus-to-cytoplasm ratio than that of the GS-rich control peptide motif (Fig. 3c). Similarly, the best performing NES-3 showed cytoplasmic enrichment of mNeonGreen3K comparable to the widely used PKIα NES, with lower nucleus-to-cytoplasm ratio than that of the control peptide motif (Fig. 3c). Representative images and line-intensity profiles for these constructs are shown in Fig. 3d. These results demonstrated that CELL-FM can generate new functional motifs with microscopy image guidance.

mNeonGreen3K-tagged peptide constructs are shown in green, mTagBFP-marked nuclei are shown in magenta, and merged images illustrate subcellular localization. Scale bars, 10 μm.

### Context-controlled virtual staining enables a unified subcellular space representation

Virtual microscopy also enables construction of a unified subcellular space representation. The concept of discrete compartments used in earlier sections, such as nucleus and cytoplasm, captures only coarse aspects of localization and cannot represent the subtleties commonly observed in real protein behavior. Prior approaches like CytoSelf^17^ attempted to learn protein-specific representations by training an image encoder with protein-ID supervision, encouraging the model to ignore cell-to-cell morphological variability. However, real microscopy images vary substantially across cells and imaging conditions, making it difficult to fully disentangle protein-specific localization from morphological noise. CELL-FM provides a generative alternative: by synthesizing fluorescence images for different proteins while holding the cellular morphology fixed, it enables virtual staining of the same cell for thousands of proteins, producing a morphology-controlled dataset in which differences across images arise primarily from protein-specific localization cues.

Using this virtual staining strategy, we fine-tuned CELL-FM on the OpenCell dataset^12^, which provides live-cell confocal images of 1,311 endogenously tagged proteins and the corresponding DNA-stained nuclear images in a uniform background of HEK293T cells (∼ 96,000 single-cell images after cropping). Then, conditioned on a shared nuclear image, we generated fluorescence images for each of the 1,311 proteins. The generated images were then embedded using a ViT image encoder and projected into a UMAP space to visualize the predicted protein localizations (Fig. 4a). Because all protein images are generated with the same morphological context, the suppression of irrelevant variations resulted in a better localization representation across multiple metrics than the representation directly computed from experimental images (Supplementary Fig. 3). This representation recovers major subcellular compartments as well as fine-grained distinctions that are difficult to label and suggestive of functional association (Fig. 4b). Within nucleolus, the UMAP embedding distinguishes the sub-compartments of fibrillar center (e.g. POLR1A and TCOF1), dense fibrillar component (FBL), and granular component (NPM3). In the nucleoplasm space, it separates functional compartments of chromatin (H2BC21), chromatin remodeling (SMARCA4), and RNA processing (SNRPF) from diffusive nucleoplasmic housekeeping enzymes (SAE1). Additionally, CELL-FM also recovers cytoplasmic compartments even though these regions are only weakly specified by the nuclear reference image. For example, the P-body proteins LSM14A, DCP1A, DDX6 and EDC4 form a tight cluster, suggesting specific interactions as previously demonstrated by CytoSelf^12,17^. These results show that virtual staining provides an effective foundation for learning a unified subcellular space representation, capturing stereotyped localization programs through virtual microscopy experiments.

**Figure 4:**
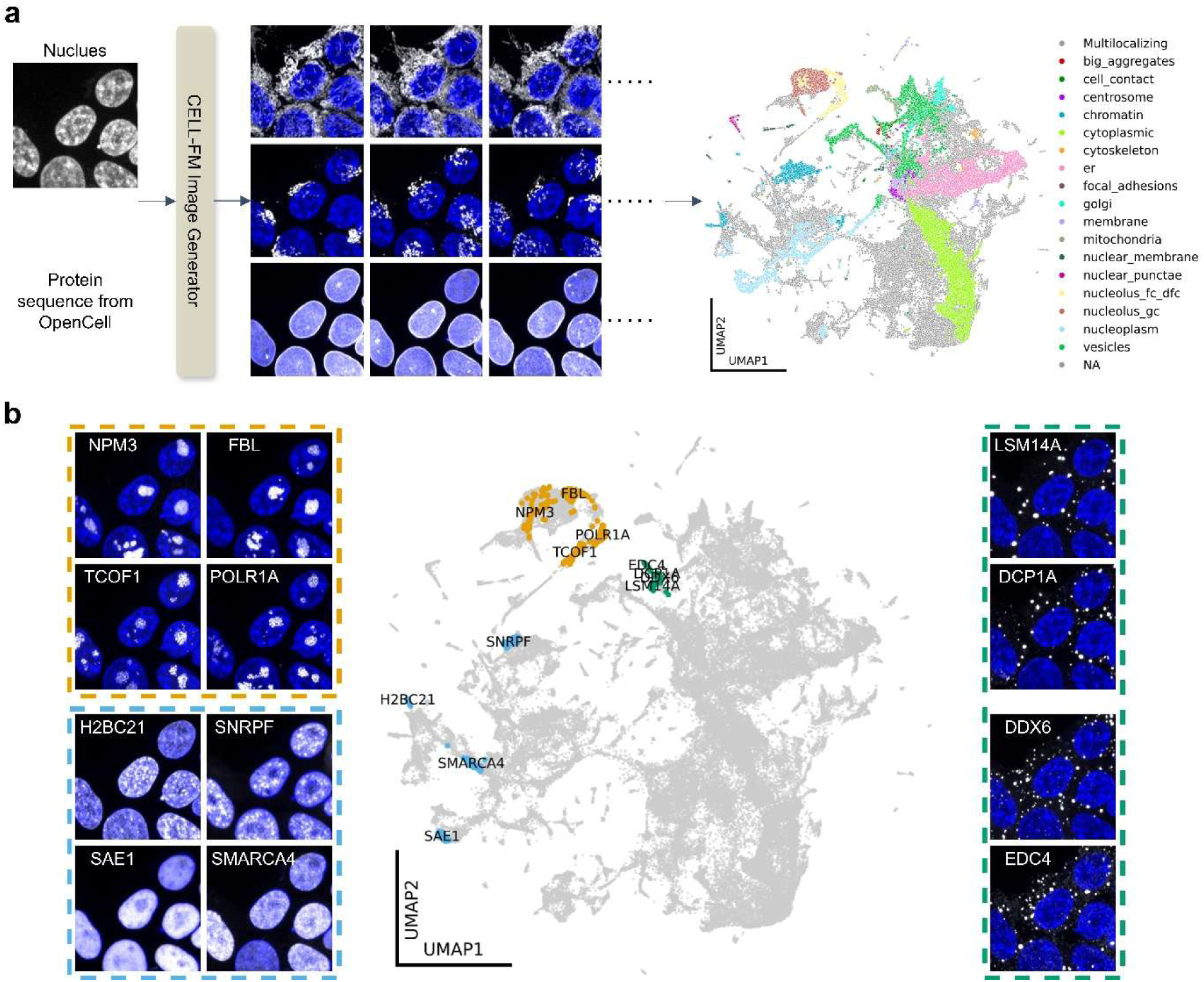
Virtual staining protein atlas generated by CELL-FM. **a.** Overview of the OpenCell virtual-staining workflow. Given a fixed nuclear morphology image and the protein sequence of each OpenCell target, CELL-FM generates the corresponding protein fluorescence image. Each generated image is then embedded with a ViT image encoder and projected into a UMAP space. **b.** Representative regions of the atlas recover fine-grained subcellular organization. Generated images separate nucleolar subcompartments, distinguish chromatin-associated patterns, and capture cytoplasmic compartments even when these structures are weakly specified by the nuclear reference image.

### CELL-FM learns sequence grammar of condensate formation

To demonstrate CELL-FM as a modeling tool for discovery and hypothesis generation, we applied it to investigating the sequence grammar of intrinsically disordered peptides (IDPs) for self-association and condensate formation through phase separation. For this purpose, we utilized the CondenSeq^24^ dataset, which is an OPS dataset covering 14,598 IDPs, each 66 a.a. long and fused to an NLS, an oligomerization domain, and a fluorescent tag, expressed in the nucleus of U2OS cells (∼ 8 million single-cell images). To quantify the condensate formation phenotype, we trained a ViT classifier to label an experimental single-cell image as 1 for condensate and 0 for no-condensate (including aberrant nucleolar and chromatin enrichment). Plotting this label against the protein expression level, as measured by the mean nuclear fluorescence intensity, can extract the saturation concentration, *c*_sat_, above which condensates are observed as small puncta in the nucleus (Fig. 5a). This saturation concentration is a key parameter to characterize phase separation behaviors^43–45^.

**Figure 5:**
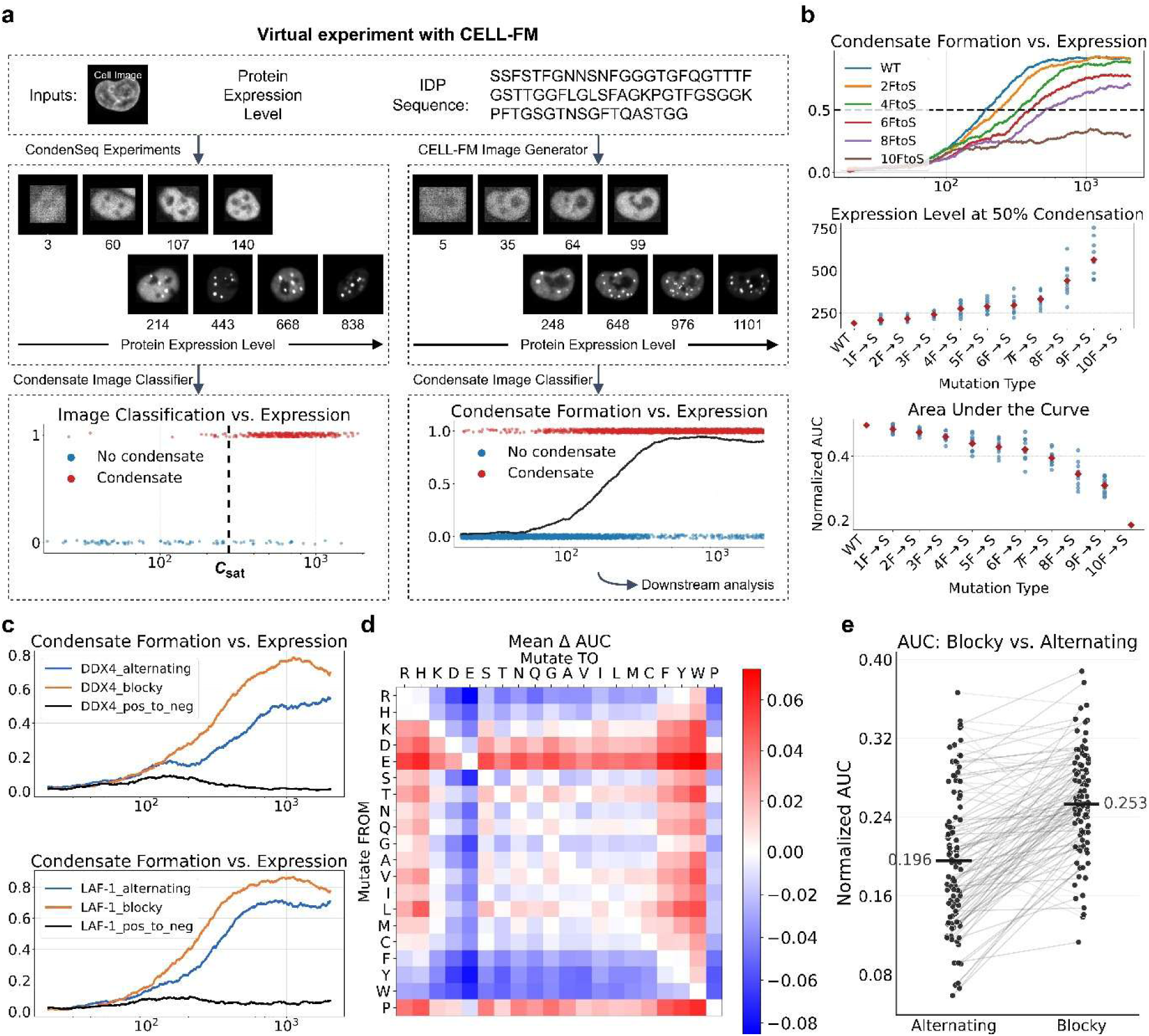
CELL-FM learns sequence grammar of nuclear condensate formation and enables virtual perturbation experiments. **a.** Virtual experiment workflow on CondenSeq. Given a fixed cell image and an IDP sequence, protein expression level is provided as a continuous conditioning variable, enabling CELL-FM to generate images for the same cellular context across an arbitrary expression range. A ViT-based condensate classifier (condensate vs no-condensate) converts each image into a binary label, which we smooth with a moving average to obtain a continuous condensate probability curve as a function of expression level. **b.** Virtual F to S mutagenesis of NUP98 IDP. Top, condensate titration curves for the wild-type sequence and mutants with increasing numbers of F-to-S substitutions. Middle, expression level required to reach 50% condensate probability. Bottom, normalized area under the condensate probability curve (AUC) for the same set of mutants. **c.** Patterning variants of natural IDPs. For DDX4 and LAF-1, engineered variants that preserve non-charged composition but alter charge patterning show enhanced condensation for blocky variants relative to alternating variants. **d.** The amino acid substitution matrix. For 100 diverse held-out IDPs, we generate images across expression levels for each substitution type and summarize the mean change in AUC relative to the unmutated sequence. **e.** Systematic test on a synthetic charged library. Across 100 paired sequences with balanced charge and fixed spacer positions, permuting only charged residues to create alternating versus blocky variants yields higher AUC for blocky sequences.

Although it is possible to train a model predicting *c*_sat_ directly from the sequence^46^, this model will not be able to capture more complicated behaviors observed in the CondenSeq dataset, many of which cannot be described by a single *c*_sat_. Therefore, we trained CELL-FM as a virtual microscopy experiment model with the CondenSeq data (with 1,058 IDPs held out for testing based on sequence similarities). Importantly, to learn the intrinsic concentration dependence of condensate formation, we explicitly introduced protein expression level as an additional, continuous conditioning variable. In this way, CELL-FM can generate microscopy images of an IDP, conditioned on a given nucleus image and at expression levels with a much higher sampling density than practically achievable in OPS experiments. Feeding the generated images to the same ViT classifier, followed by a moving average smoothing of the binary labels, provides the condensate titration curve for this IDP describing the probability of observing condensates at a given expression level (Fig. 5a).

To validate CELL-FM as an IDP condensate formation model, we performed a virtual mutation study for a 66 a.a. region from the NUP98 IDP by mutating random subsets of phenylalanine (F) into serine (S), generating condensate titration curves (Fig. 5b, top and Supplementary Fig. 4), and measuring the expression level at which the curves cross the 50% threshold as a surrogate for *c*_sat_. Consistent with the knowledge that NUP98 IDP condensate formation is driven by aromatic interactions from the FG repeats^47^, the approximate *c*_sat_ increases with the number of point mutations (Fig. 5b, middle). Moreover, when all 10 Fs are mutated, the titration curve falls below the 50% threshold for the entire tested expression level range. We also evaluated the area under the titration curve (AUC) as a threshold-free metric for condensate formation potency, which faithfully captured the same trend (Fig. 5b, bottom). As another validation, we studied two IDPs from DDX4^48,49^ and LAF-1^50^, whose condensate formation is largely charge-driven. Titration curves from CELL-FM virtual experiments again match existing knowledge on this type of IDPs: residue-shuffling mutants with blocky positive and negative charges have higher potency for condensate formation than mutants with alternating positive and negative charges, whereas composition mutants flipping all positive residues to negative are unable to form condensates (Fig. 5c).

We next scaled this analysis up for two large-scale *in silico* mutation studies. To investigate the contribution of amino acid composition, we selected 100 test set IDPs with relatively diverse amino acid composition (to reduce bias from rare residues) and mutated each type of amino acid to each of the other types. For each sequence and each substitution type, we generated 512 images across expression levels on a log scale and computed the AUC of the titration curves. We then aggregated the mutation effect as the mean change in AUC relative to the unmutated sequence to generate the full substitution matrix (Fig. 5d). The most prominent feature of this matrix indicates that aromatic residues (F, Y, and W) and arginine (R) enhance condensate formation, consistent with their known sticker roles in the sticker-spacer model for IDP phase separation^45,51,52^. Histidine (H) is also shown as a strong sticker. Negatively charged residues (D and E) are highly disfavored for condensate formation, but the positively charged lysine (K) is more tolerated, potentially because it can engage in cation-pi interactions^53^. As an interesting observation not previously reported in experimental studies, for spacer residues of similar chemical properties, having one extra carbon in the side chain subtly but consistently reduces condensate formation (comparing S to T, N to Q, and G to A). This size dependence supports the use of spacer solvation volume to describe their contribution in the sticker-spacer theory^54^.

To investigate the contribution of charge pattern, we constructed a synthetic IDP library containing only charged residues (R, K, D, E) and neutral spacers (G, S, Q, N, T). For each of 100 randomly generated backbones (66 a.a.) with fixed spacer positions and balanced charge (15 positive, 15 negative), we constructed paired alternating and blocky variants by permuting only the charged positions. Across matched pairs, CELL-FM predicts consistently higher AUC for blocky sequences (Fig. 5e), increasing on median from 0.196 (alternating) to 0.253 (blocky). When examining CondenSeq images, we also discover that a fraction of IDPs show phase re-entrant behavior, in which condensates form at intermediate expression levels but dissolve at higher levels (Supplementary Fig. 5a). This non-monotonic response cannot be handled by a model trained to predict *c*_sat_, but the same CELL-FM model can quantify phase re-entrance simply by switching the metric to the area above the titration curve (AAC) beyond the maximum point (Supplementary Fig. 5b). From our earlier *in silico* composition mutant results, we generated the substitution matrix for phase re-entrance tendency (Supplementary Fig. 5c). Compared to their roles in driving condensate formation, aromatic residues (F, Y, and W) are still favored for phase re-entrance but not arginine (R) or histidine (H), whereas hydrophobic residues (V, I, L, M and C) surprisingly stand out as strong drivers. These observations will help the hypothesis generation to study this under-appreciated phenomenon^55,56^, which likely involves both the self-association of IDPs and their interactions with other nuclear biomolecules.

## Discussion

CELL-FM demonstrates that generative modeling of experimental readouts can serve as a powerful alternative to label-based prediction based on large-scale biological screens. By learning to synthesize microscopy images from protein sequence and cellular context, as well as to propose sequences consistent with image phenotypes, it enables virtual experiments that flexibly support localization prediction, motif discovery and analysis, image representation learning, and large-scale mutation studies for hypothesis generation. The ability to generate controlled virtual staining images allows thousands of proteins to be visualized within a shared cellular context, which is nearly impossible by experiments and enables harmonized subcellular representations that capture subtle localization differences beyond discrete compartment labels. Sequence generation could lead to applications in protein engineering, where desired spatial phenotypes could guide sequence design. Lastly, as illustrated by our IDP virtual experiments using both protein sequence and expression level as controlled inputs, the unified generative framework of CELL-FM can easily cover other types of variables across distinct modalities, such as cell type labels, genetic perturbations, and descriptors of measurement conditions.

At the same time, several limitations remain. Generative models are still bound by the training data distribution and inherit biases within the training data. For example, immunofluorescence artifacts can be observed in some images generated by HPA-trained CELL-FM. Therefore, their output should be interpreted cautiously and can be problematic when the input falls outside of the training data distribution. Aligning datasets collected in different experiments, under different conditions, or from different cell types can be challenging. Our HPA-pretraining and OpenCell-finetuning strategy for virtual staining provides one practical solution, and more systematic approaches for cross-dataset harmonization will be needed for broader generalization. Finally, while CELL-FM captures static localization patterns and cell-to-cell variability, extending virtual experiments to dynamic behaviors, multi-channel interactions, or perturbation-dependent phenotypes remains an important direction for future work.

## Acknowledgements

We thank Kalli Kappel for inspiring discussions. We thank Biohub SF Scientific Compute Platform for their support. B.H. is a Biohub San Francisco Investigator.

## Author Contributions

D.Z. and B.H. conceived and designed this study. D.Z. performed all computational work. K.H. performed the validation experiments. D.Z. and B.H. interpreted the results. B.H. supervised the project and acquired funding. D.Z. and K.H. prepared the figures. D.Z., K.H., and B.H. wrote the manuscript.

## Data availability

The Human Protein Atlas data can be downloaded from the CZI virtual cell platform (https://virtualcellmodels.cziscience.com/dataset/hpa-subcellular-section).

The OpenCell data can be downloaded from the OpenCell website (https://opencell.sf.czbiohub.org/).

The CondenSeq data can be downloaded from Bioimage Archive (https://www.ebi.ac.uk/biostudies/bioimages/studies/S-BIAD1738).

## Code availability

The code is available on GitHub at https://github.com/BoHuangLab/CELL-FM.

## Supplementary Information

### Methods

#### Overview of CELL-FM

CELL-FM is a unified generative framework that connects protein sequences and fluorescence microscopy images through bidirectional conditional generation. The model represents protein fluorescence images as a continuous generative object, trained in the latent space of a pretrained VAE, and protein sequences as a discrete generative object, trained with a discrete diffusion objective. Both modalities are integrated through a shared transformer encoder that fuses protein sequence tokens derived from a pretrained protein language model and image patch tokens derived from multi-channel microscopy inputs (protein and cellular contexts).

Three modality-specific heads branch from the shared representation: an image generator trained by flow matching^28^, an image reconstruction head trained by a denoising masked autoencoder (dMAE) objective^29,30^, and a sequence generator trained by discrete diffusion^31,32^.

#### Image latent space with pretrained VAE

To efficiently model high-resolution fluorescence patterns, we generate protein images in the latent space of a pretrained VAE. We adopt the same VAE architecture used in Stable Diffusion^27^ with two downsampling stages. During VAE training, each target protein fluorescence image is encoded by the VAE encoder into a Gaussian latent distribution, sampled via the reparameterization trick^26^, and reconstructed by the VAE decoder. The encoder applies two stages of spatial downsampling, producing a compact latent with 4 channels: from 256 × 256 to 64 × 64 for HPA/OpenCell, and from 160 × 160 to 40 × 40 for CondenSeq. The VAE is trained with a standard combination of MSE reconstruction loss and a KL divergence regularization loss. To preserve fine structural detail and avoid overly smooth reconstructions, we use a small KL weight of 10^-4^ in all experiments.

#### Unified encoder

The unified encoder is a transformer backbone that processes a single token sequence formed by concatenating image tokens and protein-sequence tokens, enabling early and deep multimodal fusion via standard multi-head self-attention. Latent images are patchified into non-overlapping 4 × 4 patches and linearly projected into the encoder hidden space. Protein sequences are embedded with a pretrained protein language model (ESMC-600M^25^) and then projected to the same hidden dimension. We apply sinusoidal positional embeddings within each modality to encode spatial order for image patches and sequential order for protein tokens. We also add a learnable token-type embedding to indicate whether a token originates from the image or sequence stream. For the image generation objective, the continuous time variable *t* is encoded as a time token and concatenated to the image token sequence, allowing the encoder to condition all subsequent computation on the diffusion or flow time step. Across all datasets, the encoder uses a hidden size of 1152, a maximum protein sequence length of 2048, and multi-head self-attention with 8 heads and head dimension 64. In addition, we use 8 transformer layers for HPA/OpenCell and 4 transformer layers for CondenSeq. The encoder produces a shared latent representation that is consumed by three decoders described below.

#### Flow matching for protein image generation

Flow Matching^28^ serves as a simulation-free generative framework that provides a compelling alternative to score-based diffusion^57,58^. Instead of training a denoiser across discrete noise levels via stochastic sampling, it learns a velocity field that deterministically transforms a simple base distribution (such as Gaussian noise) into the target data distribution. This approach streamlines the training process and often results in higher fidelity generative performance.

We train the image generator using a flow matching objective in latent space. For each training sample, we sample a scalar time *t* ∼ *U*(0,1), a Gaussian noise latent *x*_0_ ∼ *N*(0, *I*), and form a linear interpolation *x*_t_ = (1 − *t*)*x*_0_ + *tx*_1_, where *x*_1_ is the VAE latent of the target protein fluorescence image. Under this linear path, the corresponding velocity is constant:

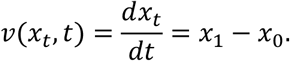

The image generator *v*_θ_(⋅) is conditioned on the unified encoder representation, which integrates cellular context *x*^cell^ and protein sequence *s*, and is trained to regress the target velocity:

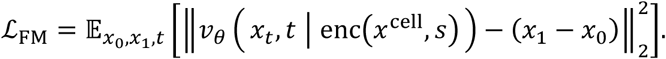

At inference time, we sample *x*_0_ ∼ *N*(0, *I*) and integrate the ODE:

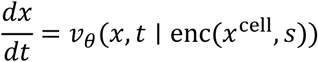

from *t* = 0 to *t* = 1 to obtain a generated latent *x̂*_1_, which is then decoded into the protein fluorescence channel using the VAE decoder.

The image generator *v*_θ_ is implemented with an MMDiT^59^ architecture and operates on non-overlapping 2 × 2 patches in the VAE latent space. We use 64 head dimensions across datasets and adjust model capacity by dataset: for HPA/OpenCell, *v*_θ_ has 8 transformer layers with 18 attention heads, whereas for CondenSeq it uses 4 transformer layers with 8 heads.

#### Discrete diffusion for sequence generation

To enable image-to-sequence generation, we train a discrete generative model over amino acid tokens using an order-agnostic autoregressive diffusion model (OA-ARDM)^31,32^. Specifically, we define a corruption process that progressively masks tokens in the sequence and a reverse model that predicts masked tokens conditioned on the unmasked subset. During training time, we sample a random masking pattern, then train the sequence head to predict the masked residues given the visible residues and the unified encoder representation.

Denoting the full sequence by *s* and the visible subset by *s*_vis_, the training objective is a masked-token negative log-likelihood:

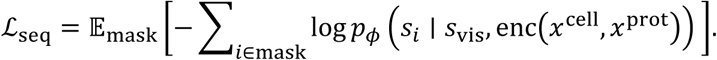

The sequence decoder is implemented as a 2-layer MLP projection head that is applied to encoder outputs producing logits over the amino acid vocabulary for each masked site.

During the generation phase, we initialize the sequence with all residues masked and condition the model on observed microscopy images. The model then iteratively unmasks and fills residues according to a randomly sampled order; at each step, it predicts a categorical distribution for the next position to be revealed, samples the corresponding amino acid token, and appends it to the visible set. This process is repeated until all positions are filled, yielding a complete sequence.

#### Denoising masked autoencoder

In addition to the two generative objectives above, we include a denoising masked autoencoder (dMAE)^29,30^ head to reconstruct masked image tokens. For the image stream fed into the unified encoder, we randomly mask a fraction *r* of non-overlapping patches and keep only the visible patches as encoder inputs. For reconstruction, we insert learnable mask tokens at the masked positions and apply a decoder to predict the missing content in VAE-latent space, i.e., the latent patch targets corresponding to the target protein fluorescence channel.

The dMAE objective encourages the unified encoder to learn robust, context-aware representations, reducing overfitting to a single modality and improving optimization during joint training (Supplementary Fig. 1b). Let *x*_1_ denote the VAE latent of the target protein fluorescence image and *x̂* the reconstructed latent predicted by the dMAE decoder. The reconstruction loss is

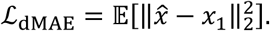

The reconstruction decoder follows the standard MAE^29^ design, and we set the masking ratio to *r* = 0.5. The decoder takes the unified encoder outputs for visible patches, inserts learned mask tokens at the masked locations and processes the resulting full patch sequence with a lightweight transformer decoder. The decoder outputs are then linearly projected back to VAE-latent patch space to predict the latent targets for all patch positions.

We use the same decoder width and attention configuration across datasets with hidden size 512, 8 attention heads with head dimension 64, and 4 transformer layers for HPA/OpenCell and 2 transformer layers for CondenSeq.

#### Joint training

Training proceeds in two stages:

##### Stage 1: representation pretraining

We jointly optimize a weighted sum of the flow matching, sequence, and dMAE objectives,

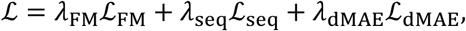

using an image masking ratio *r* = 0.5. We set *λ*_FM_ = *λ*_seq_ = *λ*_dMAE_ = 1. This stage encourages the unified encoder to learn robust multimodal representations that support both reconstruction and conditional generation.

##### Stage 2: generative finetuning

We disable masking (*r* = 0) and finetune the model using only the generative losses ℒ_FM_ and ℒ_seq_ (with *λ*_FM_ = *λ*_seq_ = 1). This stage focuses model capacity on high fidelity image and sequence generation.

Both stages are trained for 50,000 steps using the AdamW^60^ optimizer with a 1,000-step linear warmup to a peak learning rate of 3 × 10^-4^, followed by a linear decay to zero. We use a batch size of 128 for HPA and OpenCell, and 512 for CondenSeq.

#### Protein expression level conditioning for CondenSeq

To model concentration dependent condensate formation in CondenSeq, we incorporate an explicit conditioning variable *e* that represents protein expression level. For each training image, *e* is computed as the mean protein intensity within the nuclear mask minus the dataset-level mean background intensity. The resulting scalar is mapped to the encoder hidden dimension with a linear projection and appended to the image token sequence as a dedicated conditioning token before being processed by the unified encoder. Conditioning on *e* enables CELL-FM to generate protein fluorescence images at any expression level, supporting generation across a continuous expression range while fixing the sequence and cellular context.

#### Comparison to CELL-Diff

CELL-FM builds on the original CELL-Diff^23^ framework but makes several improvements in model scalability, generation fidelity, and quantitative benchmarking. Firstly, CELL-FM replaces the DDPM^57^ objective in CELL-Diff with a flow matching^28^ formulation for protein image generation in latent space, which simplifies training dynamics and improves sample quality. On the sequence side, CELL-FM updates the protein language model embedding from ESM2^61^ to ESMC^25^, providing a stronger pretrained sequence representation for multimodal fusion.

Architecturally, CELL-FM uses a transformer-based unified backbone to process and integrate both sequence tokens and image tokens, whereas CELL-Diff relied on a hybrid design combining U-Net components (for images) with transformer modules (for cross-modal fusion). Beyond model changes, CELL-FM also extends the scope of benchmarking and applications. We introduce distribution-based evaluation using a microscopy image trained embedding network for FID/KID-style comparisons between sets of generated and experimental single-cell images, and we evaluate on a larger, more comprehensive held-out protein split than in the original CELL-Diff study. Finally, while CELL-Diff primarily targets general subcellular localization prediction, CELL-FM is further extended to CondenSeq, a focused dataset for nuclear condensates. This setting enables controlled *in silico* titration by explicitly conditioning generation on protein expression level, allowing us to probe how sequence features modulate the onset, morphology, and dissolution of condensate formation.

#### Model ablation

We first performed ablations on the denoising masked autoencoder (dMAE) auxiliary loss used during representation pretraining. For dMAE, we trained two models that were identical in architecture, data, and finetuning protocol, differing only in Stage 1: one used the full multi-task objective (flow matching, sequence diffusion, and dMAE) while the other omitted dMAE; both were then finetuned in Stage 2 with masking disabled using only the generative losses (flow matching and sequence diffusion). Including dMAE in Stage 1 improves model convergence and reaches lower loss earlier (Supplementary Fig. 1b). Next, we varied the patch size used to tokenize VAE latents for image generation and found that smaller patches consistently improved fidelity (Supplementary Fig. 1c), likely because finer tokenization increases spatial bandwidth and reduces patch-boundary artifacts, albeit at higher compute and memory cost due to longer sequences. Finally, to quantify how protein diversity affects generalization to unseen proteins, we trained models on progressively larger subsets of the HPA training split (500, 1,000, 3,000, 6,000, and 11,694 proteins) and evaluated on the fixed held-out test set; performance improved monotonically with scale, with a pronounced reduction in distributional distance (FID/KID) when increasing from 1,000 to 3,000 proteins (Supplementary Fig. 1d). Notably, OpenCell contains 1,311 proteins, close to this transition point, suggesting that OpenCell alone is unlikely to provide sufficient protein diversity to train a general-purpose sequence-to-image generator without additional data or pretraining.

#### Data preprocessing and augmentation

Across all datasets, we apply random flips and random rotations for data augmentation. All fluorescence channels are normalized to the range [−1, 1].

##### HPA

We use the Human Protein Atlas (HPA)^10,11^ Subcellular Section images (v23), consisting of confocal immunofluorescence images with four channels: microtubules, endoplasmic reticulum (ER), nuclei, and protein. To train CELL-FM, we start with the SubCell-processed HPA crops^18^, in which cells are segmented and exported as 1024 × 1024 single-cell images centered on each detected cell. We then resize each crop to 256 × 256 for model input. Protein sequences are retrieved from UniProt and mapped to images via gene identifiers.

##### OpenCell

We use OpenCell^12^ microscopy images of live HEK293T cells, where each target protein is endogenously tagged with split-mNeonGreen2 and DNA is stained with Hoechst 33342. For training, we use maximum-intensity projection images and generate 256 × 256 single-cell crops centered on nuclei detected from the Hoechst channel. Protein sequences are retrieved from UniProt and mapped to the corresponding tagged protein targets.

##### CondenSeq

CondenSeq^24^ is a pooled imaging dataset with an *in situ* sequencing assay that quantifies nuclear condensate formation across large libraries of short synthetic sequences. Cells are imaged in the Hoechst and GFP/SNAP-tag channels for phenotype readout. The original study reports phenotype imaging on an Opera Phenix system in confocal mode, with images acquired at a single z-plane. The assayed sequences are fixed length (66 amino acids). We use the GFP phenotype images together with the accompanying sequence library. To standardize image geometry, we zero-pad or crop images to 160 × 160 while keeping the nucleus centered.

#### ViT model for image embedding

To evaluate generated protein images with distributional metrics (msFID/msKID), we require an embedding space that is tuned to fluorescence microscopy rather than natural images. We therefore train a microscopy-specific image encoder based on a ViT using protein-ID classification as a pretext task (Supplementary Fig. 2b). Concretely, we concatenate the cellular context with the target protein image and split the single-cell crop into non-overlapping 4 × 4 patches. Each patch is flattened and linearly projected into a token embedding, and the token sequence is processed by a ViT encoder with 12 transformer blocks (8 attention heads per block) and an MLP sublayer of width 2,048. We then apply average pooling over the output tokens to obtain a fixed-length feature vector (512 dimensions), which is fed to a linear classification head to predict protein identity during training. After pretraining, we discard the classification head and use the pooled ViT representation as the latent image embedding for downstream comparisons.

Pixel-wise metrics such as PCC and PSNR tend to reward mean-like predictions that blur or suppress localization structure (Supplementary Fig. 2a). In contrast, embedding-based distances compare distributions of images and are sensitive to higher-level phenotypic attributes. In Supplementary Fig. 2c, the left example shows substantial overlap between predicted and experimental samples in embedding space, resulting in a low msFID (1.55) consistent with matched punctate organization. The right example exhibits clear separation between the two distributions and a correspondingly high msFID (83.16), indicating a failure to reproduce the correct phenotype.

Finally, Supplementary Fig. 2d visualizes the learned embedding of the HPA held-out test set using UMAP, colored by localization labels. The embedding organizes proteins by subcellular phenotype, with strong separation between nuclear and cytoplasmic classes and coherent grouping of major compartments, supporting the use of these ViT features for FID/KID-style evaluation.

#### Computation of msFID/msKID

Let 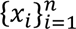 and 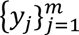 denote ViT embeddings of generated and experimental images, respectively, where *x*_i_, *y*_j_ ∈ ℝ^d^ and *d* = 512.

##### Microscopy-adapted Fréchet Inception Distance (msFID)

FID^33^ approximates each embedding set as a multivariate Gaussian and computes the Fréchet distance between the two Gaussians. Define empirical means and covariances

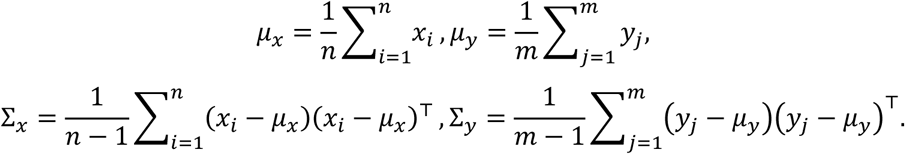

Then

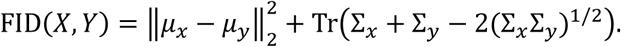

##### Microscopy-adapted Kernel Inception Distance (msKID)

KID^34^ is the squared Maximum Mean Discrepancy^62^ (MMD) between two embedding distributions using a polynomial kernel. We adopt the common third-degree polynomial kernel:

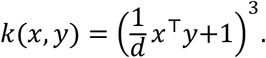

The unbiased KID estimator is

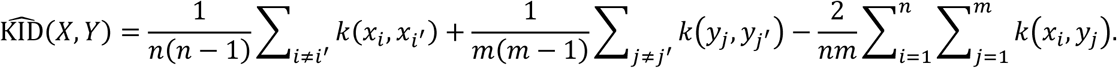

However, the available data for each protein is constrained, typically ranging from dozens to a few hundred single-cell cropped experimental images. Given this limited sample size, the unbiased estimator exhibits high variance and frequently yields non-physical negative values. To ensure numerical stability and strict non-negativity, we employ the biased estimator (V-statistic)

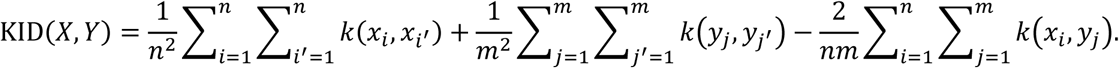

While this introduces a positive bias of order *O*(1/*N*), it provides a more robust signal for comparison in this low-data regime.

#### Image-conditioned generation of NLS and NES motifs

For motif generation, we fine-tuned the HPA-pretrained CELL-FM model on the complete HPA dataset. This model variant operates at an input resolution of 512 × 512, with a patch size of 8 for the image encoder and a patch size of 4 for the image generator operating on the corresponding VAE latent representation. Because the model was trained on the complete dataset, it was used for sequence generation only. Sequence generation was conditioned on one experimental single-cell image selected to represent each target phenotype (Fig. 3a), together with the corresponding cellular-context channels for the nucleus, ER, and microtubules. Candidate motifs were generated through masked infilling at the C terminus of a 24-residue scaffold, MPSQGSLGAAPPEVAPDSSETEEG, which provided the sequence decoder with a constant and localization-neutral sequence context. For each target motif length, the scaffold was extended with the corresponding number of <MASK> tokens. These positions were then filled using order-agnostic autoregressive diffusion. Specifically, the masked positions were randomly permuted, and one position was filled at each step. The model predicted a categorical distribution over amino acids at the selected position, conditioned on the input images and all residues generated in previous steps. An amino acid was sampled from this distribution and inserted into the sequence before proceeding to the next masked position. We generated motifs ranging from 10 to 25 amino acids in length, with 20 sequences sampled at each length for each phenotype. The resulting 320 sequences per phenotype were used for the compositional analysis shown in Fig. 3b. For experimental validation, we independently generated 40 sequences at each length, ranging from 5 to 25 amino acids. To prioritize candidates for experimental testing, each generated motif was modeled in complex with its cognate nuclear-transport receptor using AlphaFold3^41^. NLS candidates were modeled with the import receptor KPNA1, whereas NES candidates were modeled with the export receptor CRM1. We selected the first 10 NLS and 10 NES candidates with an interface predicted template modeling score (ipTM) greater than 0.8 for experimental validation.

#### Preparation of plasmids for NLS/NES validation

All plasmids used for transient overexpression of generated peptide motif-fused mNeonGreen3K were cloned by In-Fusion cloning as manufacturer’s instruction (TaKaRa). Each peptide coding DNA was amplified by PCR with two primers, and plasmid backbone was also linearized by PCR with pHaloTag7-mNG3K-IRES-NLS-mTagBFP-NLSx2 plasmid. Cloned plasmids were transformed into HST08 Stellar Competent cells (TaKaRa), and all sequences of purified plasmids were verified by nanopore sequencing (Plasmidsaurus).

#### Transient overexpression of NLS/NES-fused mNeonGreen3K in mammalian cells

10,000 U2OS wild-type (ATCC) cells were seeded in each well in 96-well plates (CellVis), then incubated overnight at 37 °C, 5 % CO2 in a humidified incubator (Panasonic). 200 ng of each plasmid was transfected to cells in each well with JetOptimus (Polyplus) as manufacturer’s protocol. After 3 hours of transfection, media was replaced with fresh McCoy’s 5A medium (Gibco) supplemented with 10 % FBS (UCSF cell culture facility), 1x Penicillin/Streptomycin, 1 mM sodium pyruvate, and 1x GlutaMax (Gibco), then incubated overnight at 37 °C 5 % CO2 for overexpression of proteins.

#### Live-cell imaging and image analysis

Live cell imaging of transiently transfected U2OS cells in imaging medium (FluoroBrite + 1x GlutaMax + 1x Pen/Strep) was acquired using a CSU-W1 spinning disk LFOV (Large Field of View) high-speed confocal microscope, with a Plan Apo λ 60x/NA 1.4 oil immersion objective lens and Zyla 4.2 sCMOS (Andor) for 2-channel imaging on an inverted Eclipse Ti (Nikon) microscope body (Nikon Imaging Center, UCSF). For 2-channel (NLS-mTagBFP-NLSx2 for nucleus and peptide-mNeonGreen3K for subcellular localization validation) imaging, 405 nm and 488 nm lasers (Voltran) were used to excite mTagBFP and mNeonGreen3K for fluorescence signal in cells. Single z-slice of mTagBFP, and 13 z-slices of mNeonGreen fluorescence images were taken with 0.5 μm steps, ranging 6 μm from -3 μm to 3 μm relative to z-axis focal length adjusted by perfect focus system (PFS, Nikon).

Images of mNeonGreen3K were processed with maximum intensity projection. For image analysis, automatic cell and nucleus segmentation were applied to batch analysis of images. mNeonGreen3K signals were first used for cell segmentation, then mTagBFP signals were used as a reference channel for nucleus segmentation to each single segmented cell. For nucleus segmentation, we trained a custom CellPose v3^63^ model with supervised training on more than 50 cells. All experiments were conducted in biological triplicates.

#### Comparison of experimental and virtual-staining protein localization representations

We compare the organization of protein localization representations constructed from experimental OpenCell images and CELL-FM virtual-staining images. Both image sets were embedded using the same ViT image encoder and projected into UMAP space, with points colored by annotated OpenCell subcellular location classes (Supplementary Fig. 3a). Compared with the experimental atlas, the virtual-staining atlas shows tighter and better separated localization-associated regions, suggesting that fixing the cellular context reduces nuisance variation from differences in cell morphology, nuclear state, and imaging field.

We quantified this effect by applying k-means clustering in the embedding space and comparing the resulting clusters with OpenCell localization labels (Supplementary Fig. 3b). Virtual-staining embeddings improved all evaluated clustering metrics, including Adjusted Rand Index (0.384 to 0.440), V-measure (0.615 to 0.673), Fowlkes-Mallows score (0.464 to 0.516), and Hungarian matching accuracy (0.425 to 0.497). B^3^ scores also increased, with precision improving from 0.683 to 0.720, recall from 0.358 to 0.433, and F1 from 0.470 to 0.541. These results indicate that context-controlled virtual staining yields a more localization-discriminative embedding space by suppressing variation unrelated to protein identity, thereby improving protein localization clustering and downstream atlas analysis.

#### ViT model for condensate image classification

To convert generated CondenSeq phenotype images into a standardized, quantitative readout of phase separation, we trained a supervised image classifier that predicts whether the IDP in the nucleus contains condensate puncta. The original CondenSeq annotated four phenotypic categories: nucleolar, chromatin, other-condensate, and no-condensate. For our virtual experiments, we collapse these into a binary task by treating other-condensate as the positive class (condensate) and grouping nucleolar, chromatin, and no-condensate as the negative class (no-condensate). To collect the training and testing data, we manually curated 10,056 single-cell images, consisting of 8,442 negative examples (no-condensate) and 1,614 positive examples (condensate). To evaluate generalization under a balanced test distribution, we held out 1,000 images for testing (500 condensate and 500 no-condensate) and used the remaining images for training. We use a ViT model that takes single-cell crops patchified into non-overlapping 4 × 4 patches as input. Each patch is flattened and linearly projected into a 512-dimensional token embedding. The token sequence is processed by an 8-layer transformer encoder with 8 attention heads per layer and an MLP width of 2,048. On the balanced held-out test set, the classifier achieves 92.6% accuracy with macro precision 0.933, macro recall 0.926, and macro F1 0.926. During virtual experiments, we apply the classifier to each generated image to obtain a binary label (0 for no-condensate, 1 for condensate). We then smooth these discrete outcomes with a moving average across neighboring expression levels to obtain a continuous condensate probability curve. The moving average window size is set to one-eighth of the sampled protein expression range. This curve serves as an *in silico* analogue of experimental concentration titration, enabling direct comparisons of how sequence mutations or charge patterning shift the onset and strength of condensation.

#### Computation of area under the curve (AUC) and area above the curve (AAC)

To quantify the condensate-forming propensity of a protein, we use the area under the condensate probability curve (AUC) as the metric. For a given sequence, we first uniformly sample protein expression levels in log space over the range 20 to 2048, matching the expression level range of the experimental CondenSeq data. At each sampled expression level, we generate a fluorescence image using the CELL-FM image generator and classify the image with the pretrained condensate image classifier. This produces a binary condensate label for each expression level. We then smooth these discrete predictions using a moving average to obtain a condensate probability curve as a function of expression level. Let *p*(*c*) denote the condensate probability at expression level *c*. We define the AUC in log-expression space as

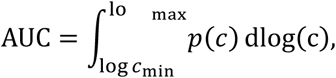

where *c*_min_ = 20 and *c*_max_ = 2048, and the normalized AUC equals to 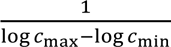 AUC. In practice, this integral is approximated numerically from the sampled expression levels. Since the AUC aggregates condensate probability across the full expression range, it provides a threshold-free measure of condensate formation propensity and avoids the need to define a condensation threshold.

To further characterize re-entrant phase behavior, we also defined the area above the condensate probability curve (AAC). This quantity measures the extent to which condensate probability decreases after reaching its maximum at higher expression levels. Using the same notation as above, we defined the AAC as

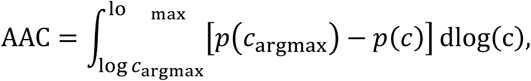

where 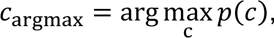 and *c*_max_ = 2048, and the normalized AAC is 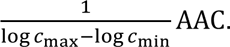

Therefore, AAC measures the decline in condensate probability after the peak of the condensate probability curve, with larger values indicating stronger re-entrant behavior.

### Sequences of the test IDPs

Residues subjected to mutation are underlined.

NUP98 IDP:

KS<u>F</u>GTP<u>F</u>GGGTGG<u>F</u>GTTST<u>F</u>GQNTG<u>F</u>GTTSGGA<u>F</u>GTSA<u>F</u>GSSNNTGGL<u>F</u>GNSQTKPGGL<u>F</u>G TSS<u>F</u>S

DDX4_blocky:

<u>RR</u>NPT<u>K</u>N<u>R</u>GFS<u>RR</u>GGY<u>RR</u>GNNS<u>R</u>ASGPY<u>RR</u>GGEGSF<u>D</u>GC<u>D</u>GGFGLGSPNN<u>E</u>L<u>D</u>P<u>DD</u>CMQ <u>E</u>TGGLFG

DDX4_alternating:

<u>RD</u>NPT<u>R</u>N<u>D</u>GFS<u>KE</u>GGY<u>RE</u>GNNS<u>R</u>ASGPY<u>DR</u>GGEGSF<u>R</u>GC<u>D</u>GGFGLGSPNN<u>R</u>L<u>D</u>P<u>RR</u>CMQ <u>R</u>TGGLFG

DDX4_pos_to_neg:

<u>ED</u>NPT<u>E</u>N<u>E</u>GFS<u>EE</u>GGY<u>ED</u>GNNS<u>E</u>ASGPY<u>EE</u>GGEGSF<u>E</u>GC<u>E</u>GGFGLGSPNN<u>D</u>L<u>D</u>P<u>DE</u>CMQ<u>E</u> TGGLFG

LAF-1_blocky:

<u>R</u>WL<u>R</u>GMSG<u>R</u>M<u>R</u>SGGGY<u>R</u>G<u>R</u>GG<u>R</u>GNGQ<u>R</u>FGG<u>DE</u>H<u>D</u>YQGGSGNGGGGNGGGGGFGGGG Q<u>D</u>SGGGGGFQ

LAF-1_alternating:

<u>R</u>WL<u>D</u>GMSG<u>R</u>M<u>D</u>SGGGY<u>R</u>G<u>E</u>GG<u>R</u>GNGQ<u>D</u>FGG<u>RR</u>H<u>R</u>YQGGSGNGGGGNGGGGGFGGGG Q<u>R</u>SGGGGGFQ

LAF-1_pos_to_neg:

<u>D</u>WL<u>E</u>GMSG<u>D</u>M<u>E</u>SGGGY<u>E</u>G<u>E</u>GG<u>E</u>GNGQ<u>E</u>FGG<u>EDEE</u>YQGGSGNGGGGNGGGGGFGGGGQ <u>E</u>SGGGGGFQ

## Supplementary Figures

**Supplementary Figure 1:**
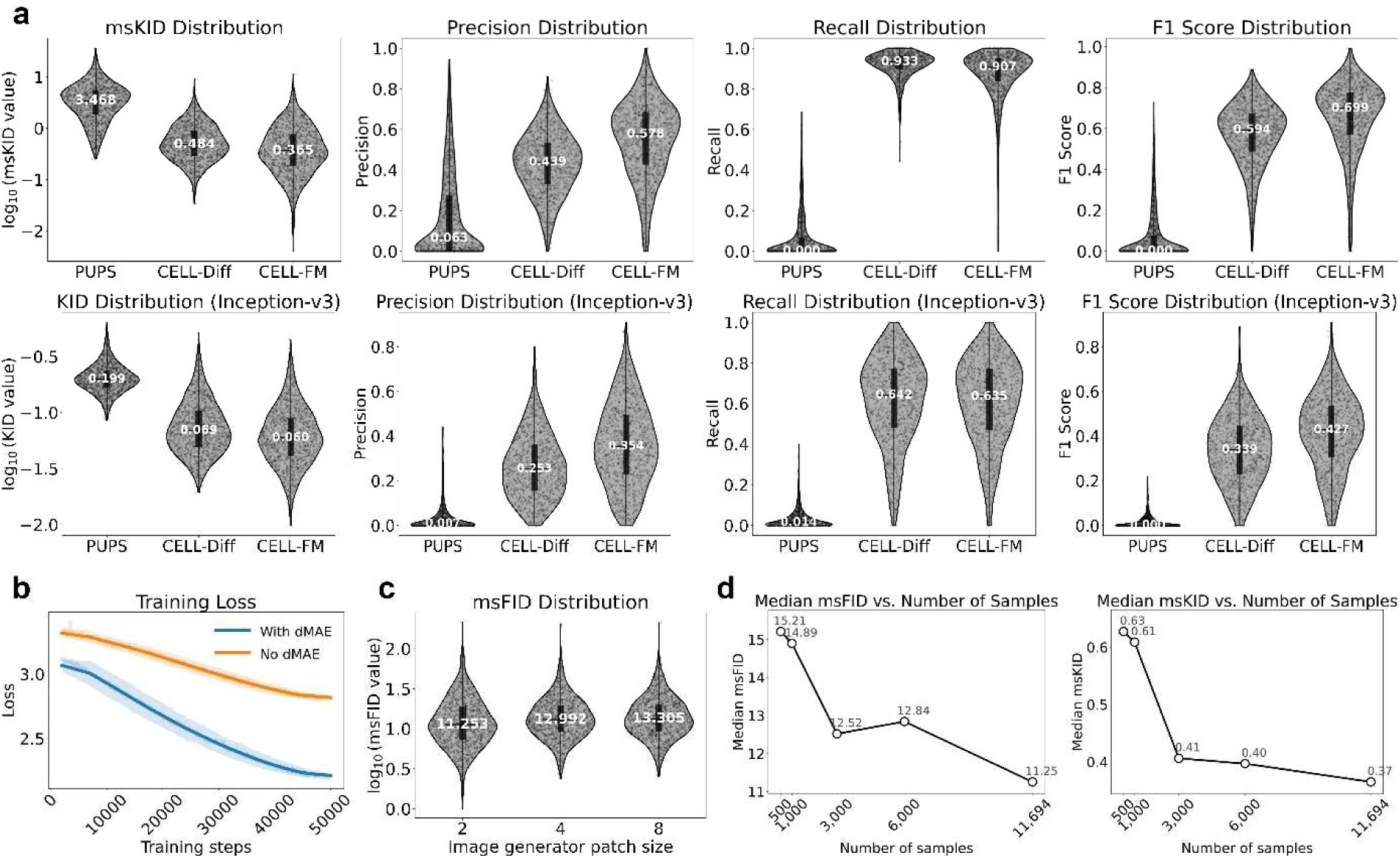
Ablation studies and evaluation controls for CELL-FM. **a.** Precision, Recall, and F1 score results on the HPA test set. **b.** Effect of the dMAE loss during generative finetuning. Training curves compare models trained with the full multi-task objective versus omitting dMAE, showing faster convergence and improved optimization stability when dMAE is included in Stage 1. **c.** Patch size ablation for image generator. Models trained with different patch sizes are evaluated, showing improved distributional fidelity with finer tokenization. **d.** Data scaling study on protein diversity. Models trained on progressively larger subsets of the HPA training proteins (with the same test set) show monotonic improvement in distributional similarity.

**Supplementary Figure 2:**
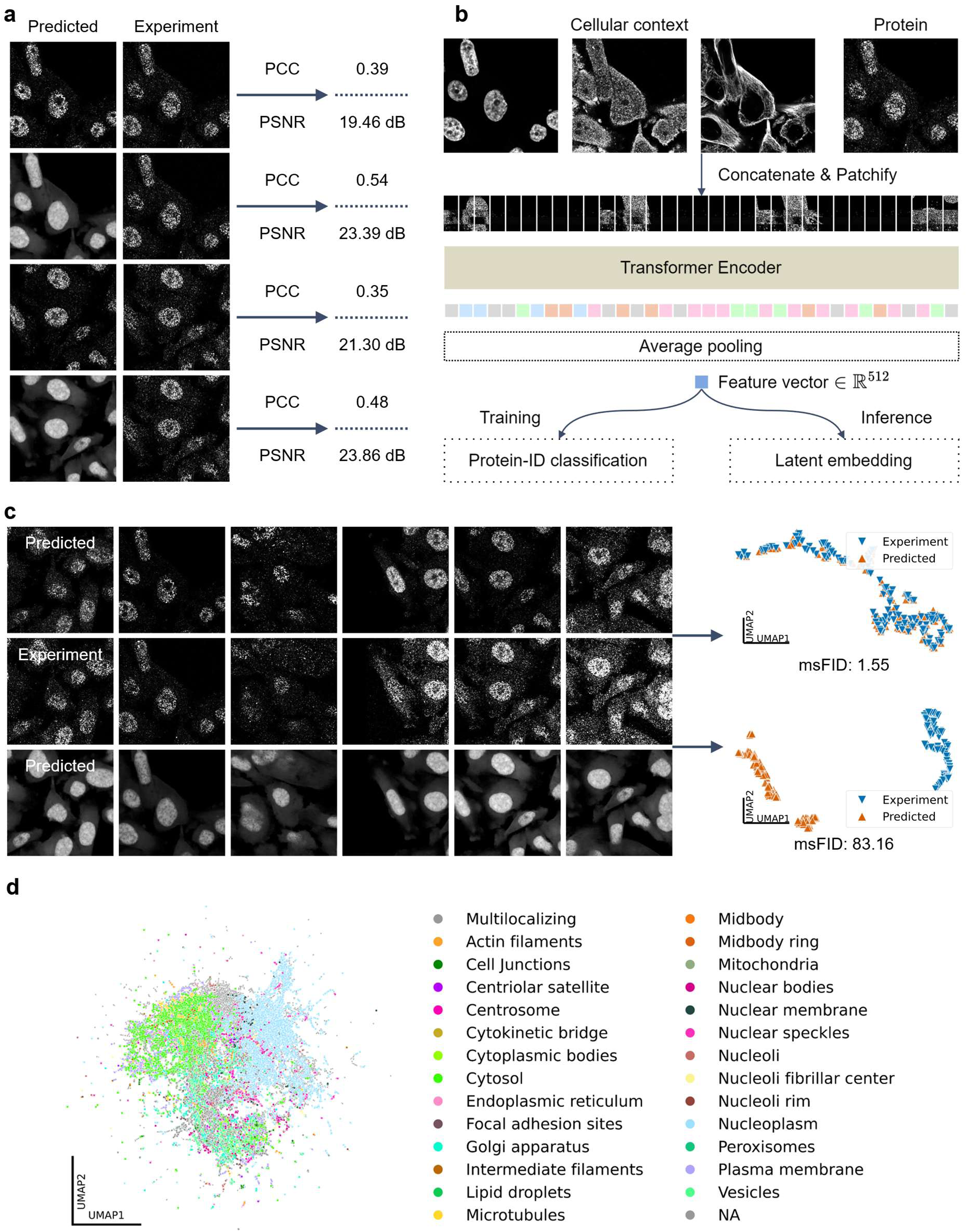
ViT embedding for distributional evaluation of generated images. **a.** Motivation for distribution-based evaluation. Pixel-wise metrics (e.g., PCC and PSNR) can favor mean-like predictions that blur or suppress localization structure, motivating distributional distances computed in a learned microscopy feature space. **b.** ViT encoder used to embed single-cell microscopy images for FID/KID-style evaluation. Cellular context channels are concatenated with the target protein channel, patchified into non-overlapping patches, projected to tokens, and processed by a ViT trained with a protein-ID classification objective on the HPA training split. After training, the classifier head is removed and pooled ViT features are used as image embeddings. **c.** Example of embedding-space comparisons illustrating low versus high distributional mismatch. Experimental and generated single-cell images are embedded with the ViT and compared as two feature distributions; substantial overlap corresponds to low msFID/msKID, whereas clear separation indicates failure to reproduce the correct phenotype. **d.** UMAP visualization of ViT embeddings for the HPA test proteins, colored by localization labels, showing that the learned feature space organizes proteins by subcellular phenotype such as coherent separation of major nuclear and cytoplasmic classes.

**Supplementary Figure 3:**
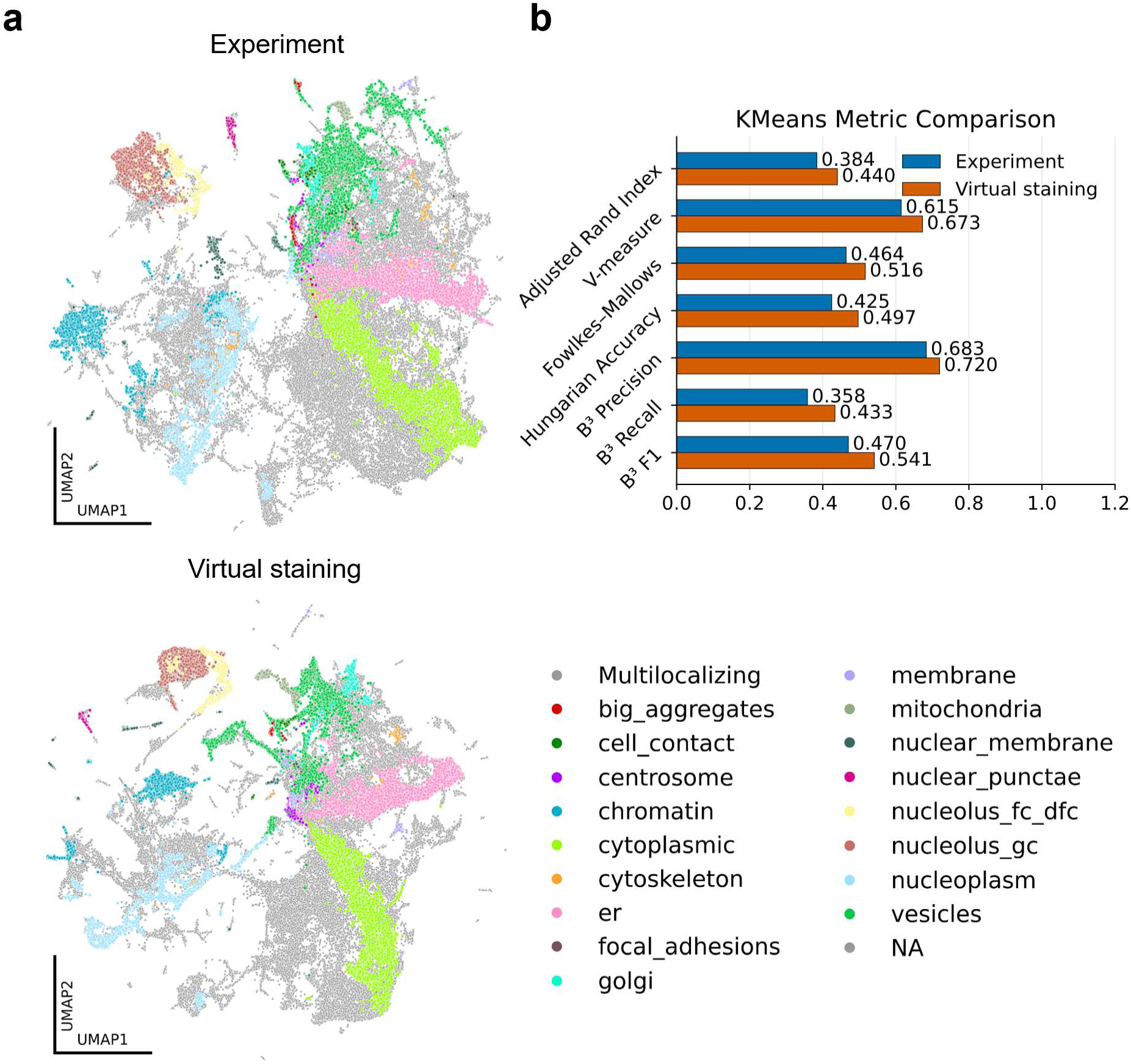
Comparison of experimental and virtual-staining protein localization atlases. **a.** UMAP visualization of image embeddings computed from experimental images and from virtual staining images, colored by annotated OpenCell subcellular location classes. **b.** Quantitative comparison of k-means clustering performance in embedding space against OpenCell location labels. Virtual staining improves multiple clustering metrics, including Adjusted Rand Index, V-measure, Fowlkes-Mallows score, Hungarian matching accuracy, and B^3^ precision, recall, and F1.

**Supplementary Figure 4:**
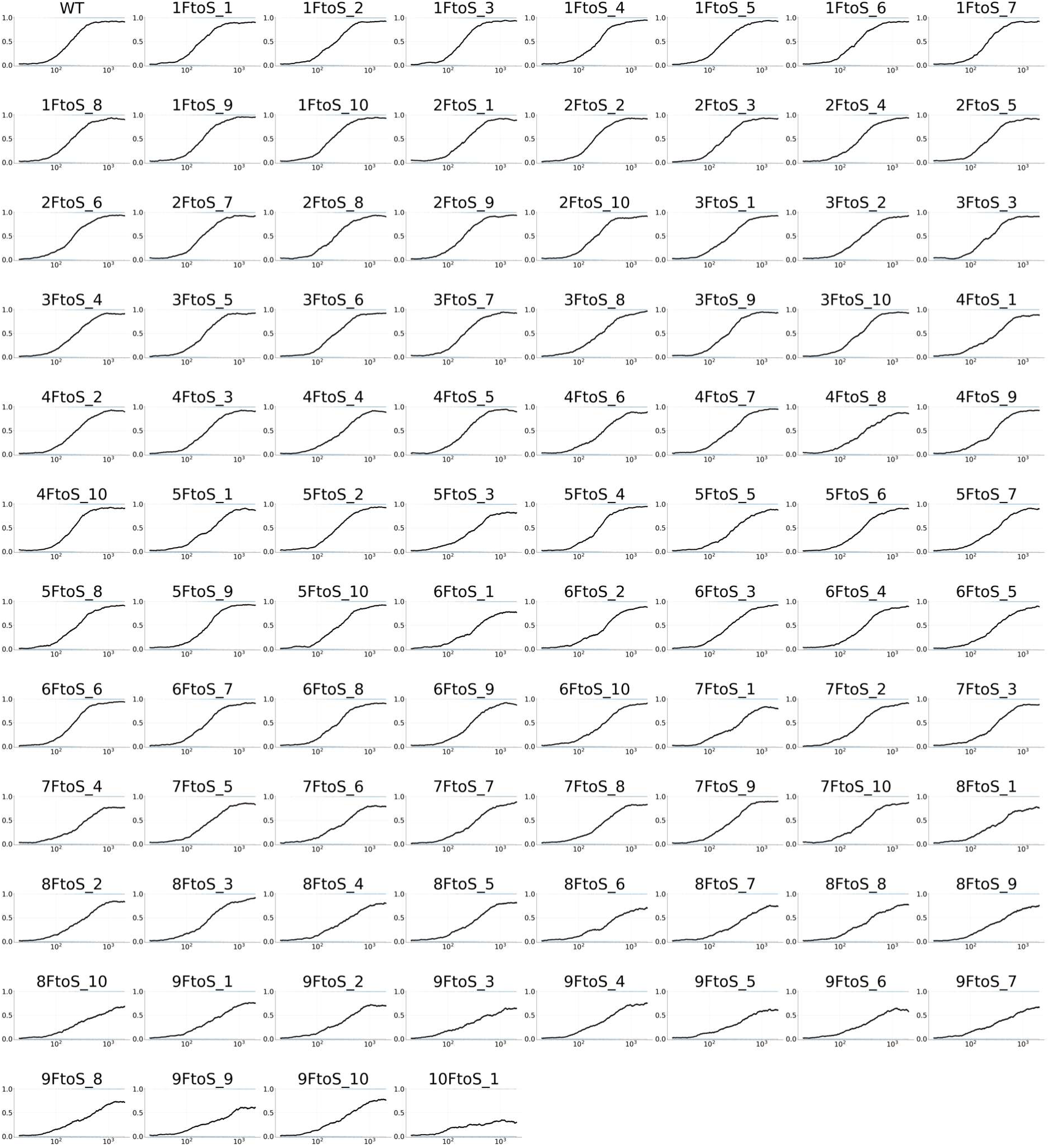
Condensate titration curves for all F to S mutations in NUP98 IDP.

**Supplementary Figure 5:**
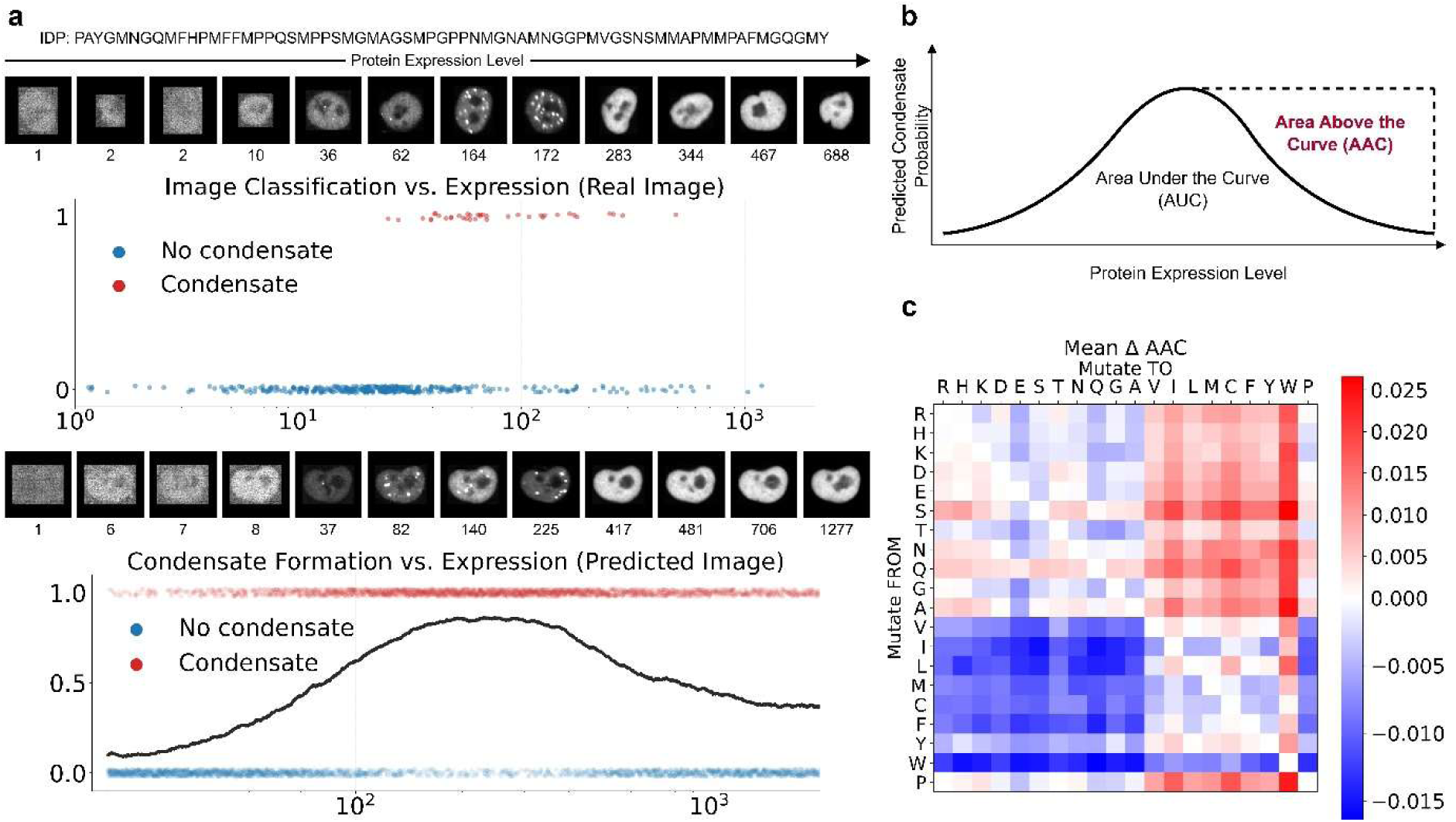
Phase re-entrant behavior on the CondenSeq dataset. **a.** Representative real and generated images for the held-out IDP sequence IDP-8991 across a range of protein expression levels. In the real images, condensates emerge at intermediate expression levels and then diminish at higher expression levels. The generated images recapitulate the same non-monotonic trend. The corresponding condensate classification plots and condensate probability curve further show that condensate formation is maximal at intermediate expression levels and decreases at higher expression levels. **b.** Schematic illustrating the area above the condensate probability curve (AAC). Larger AAC values indicate a stronger decrease in condensate propensity at high expression levels after an initial rise. **c.** Amino acid substitution effects on AAC. For 100 diverse held-out IDPs, we generated images across expression levels for every single-residue substitution type and computed the change in AAC relative to the unmutated sequence. The heatmap summarizes the mean change in AAC for each mutation pair, revealing systematic effects of amino acid identity on phase re-entrant behavior.

## References

1. Ashburner, M. et al. Gene Ontology: tool for the unification of biology. Nat. Genet. 25, 25–29 (2000).

2. Kulmanov, M., Khan, M. A. & Hoehndorf, R. DeepGO: predicting protein functions from sequence and interactions using a deep ontology-aware classifier. Bioinformatics 34, 660–668 (2018).

3. Radivojac, P. et al. A large-scale evaluation of computational protein function prediction. Nat. Methods 10, 221–227 (2013).

4. Kiselev, V. Y., Yiu, A. & Hemberg, M. scmap: projection of single-cell RNA-seq data across data sets. Nat. Methods 15, 359–362 (2018).

5. Pliner, H. A., Shendure, J. & Trapnell, C. Supervised classification enables rapid annotation of cell atlases. Nat. Methods 16, 983–986 (2019).

6. Lotfollahi, M. et al. Mapping single-cell data to reference atlases by transfer learning. Nat. Biotechnol. 40, 121–130 (2022).

7. Almagro Armenteros, J. J., Sønderby, C. K., Sønderby, S. K., Nielsen, H. & Winther, O. DeepLoc: prediction of protein subcellular localization using deep learning. Bioinformatics 33, 3387–3395 (2017).

8. Wefers, Z., Gupta, A., Ahmed, N., Zhang, X. & Lundberg, E. A comprehensive benchmark of sequence-based subcellular localization predictors for human proteins. Nat. Methods 23, 1458–1469 (2026).

9. Thumuluri, V., Almagro Armenteros, J. J., Johansen, A. R., Nielsen, H. & Winther, O. DeepLoc 2.0: multi-label subcellular localization prediction using protein language models. Nucleic Acids Res. 50, W228–W234 (2022).

10. Digre, A. & Lindskog, C. The Human Protein Atlas—Spatial localization of the human proteome in health and disease. Protein Sci. 30, 218–233 (2021).

11. Thul, P. J. et al. A subcellular map of the human proteome. Science 356, eaal3321 (2017).

12. Cho, N. H. et al. OpenCell: Endogenous tagging for the cartography of human cellular organization. Science 375, eabi6983 (2022).

13. Yang, X. et al. A public genome-scale lentiviral expression library of human ORFs. Nat. Methods 8, 659–661 (2011).

14. Matsuyama, A. et al. ORFeome cloning and global analysis of protein localization in the fission yeast Schizosaccharomyces pombe. Nat. Biotechnol. 24, 841–847 (2006).

15. Liu, C. et al. A multimodal perturbation atlas defines the phenotypic resolution of cellular morphology. bioRxiv 2026.06.01.728087 (2026) doi:10.64898/2026.06.01.728087.

16. Feldman, D. et al. Optical Pooled Screens in Human Cells. Cell 179, 787–799.e17 (2019).

17. Kobayashi, H., Cheveralls, K. C., Leonetti, M. D. & Royer, L. A. Self-supervised deep learning encodes high-resolution features of protein subcellular localization. Nat. Methods 19, 995–1003 (2022).

18. Gupta, A. et al. SubCell: Vision foundation models for microscopy capture single-cell biology. bioRxiv 10.1101/2024.12.06.627299 (2024) doi:10.1101/2024.12.06.627299.

19. Moutakanni, T. et al. Cell-DINO: Self-supervised image-based embeddings for cell fluorescent microscopy. PLOS Comput. Biol. 21, e1013828 (2026).

20. Khwaja, E., Song, Y. S. & Huang, B. CELL-E: A Text-to-Image Transformer for Protein Image Prediction. in Research in Computational Molecular Biology (ed. Ma, J.) 185–200 (Springer Nature Switzerland, Cham, 2024).

21. Khwaja, E., Song, Y., Agarunov, A. & Huang, B. CELLE-2: Translating Proteins to Pictures and Back with a Bidirectional Text-to-Image Transformer. in Advances in Neural Information Processing Systems (eds Oh, A. et al.) vol. 36 4899–4914 (Curran Associates, Inc., 2023).

22. Zhang, X., Tseo, Y., Bai, Y., Chen, F. & Uhler, C. Prediction of protein subcellular localization in single cells. Nat. Methods 22, 1265–1275 (2025).

23. Zheng, D. & Huang, B. Bridging Protein Sequences and Microscopy Images with Unified Diffusion Models. in Forty-second International Conference on Machine Learning (2025).

24. Kappel, K. et al. Characterizing protein sequence determinants of nuclear condensates by high-throughput pooled imaging with CondenSeq. Nat. Methods 22, 1464–1475 (2025).

25. Candido, S. et al. Language Modeling Materializes a World Model of Protein Biology. bioRxiv 10.64898/2026.06.03.729735 (2026) doi:10.64898/2026.06.03.729735.

26. Kingma, D. P. & Welling, M. Auto-Encoding Variational Bayes. in 2nd International Conference on Learning Representations, ICLR 2014, Banff, AB, Canada, April 14-1C, 2014, Conference Track Proceedings (eds Bengio, Y. & LeCun, Y.) (2014).

27. Rombach, R., Blattmann, A., Lorenz, D., Esser, P. & Ommer, B. High-Resolution Image Synthesis with Latent Diffusion Models. in 2022 IEEE/CVF Conference on Computer Vision and Pattern Recognition (CVPR) 10674–10685 (2022). doi:10.1109/CVPR52688.2022.01042.

28. Lipman, Y., Chen, R. T. Q., Ben-Hamu, H., Nickel, M. & Le, M. Flow Matching for Generative Modeling. in The Eleventh International Conference on Learning Representations (2023).

29. He, K. et al. Masked Autoencoders Are Scalable Vision Learners. in 2022 IEEE/CVF Conference on Computer Vision and Pattern Recognition (CVPR) 15979–15988 (2022). doi:10.1109/CVPR52688.2022.01553.

30. Wu, Q., et al. Denoising Masked Autoencoders Help Robust Classification. in The Eleventh International Conference on Learning Representations (2023).

31. Hoogeboom, E., et al. Autoregressive Diffusion Models. in International Conference on Learning Representations (2022).

32. Alamdari, S. et al. Protein generation with evolutionary diffusion. in NeurIPS 2023 Generative AI and Biology (GenBio) Workshop (2023).

33. Heusel, M., Ramsauer, H., Unterthiner, T., Nessler, B. & Hochreiter, S. GANs Trained by a Two Time-Scale Update Rule Converge to a Local Nash Equilibrium. in Advances in Neural Information Processing Systems (eds Guyon, I. et al.) vol. 30 (Curran Associates, Inc., 2017).

34. Bińkowski, M., Sutherland, D. J., Arbel, M. & Gretton, A. Demystifying MMD GANs. In International Conference on Learning Representations (2018).

35. Szegedy, C., Vanhoucke, V., Ioffe, S., Shlens, J. & Wojna, Z. Rethinking the Inception Architecture for Computer Vision. in 201C IEEE Conference on Computer Vision and Pattern Recognition (CVPR) 2818–2826 (2016). doi:10.1109/CVPR.2016.308.

36. Dosovitskiy, A., et al. An Image is Worth 16x16 Words: Transformers for Image Recognition at Scale. in International Conference on Learning Representations (2021).

37. Rowland, R. R. R. & Yoo, D. Nucleolar-cytoplasmic shuttling of PRRSV nucleocapsid protein: a simple case of molecular mimicry or the complex regulation by nuclear import, nucleolar localization and nuclear export signal sequences. Virus Res. 95, 23–33 (2003).

38. Dong, X. et al. Structural basis for leucine-rich nuclear export signal recognition by CRM1. Nature 458, 1136–1141 (2009).

39. Güttler, T. et al. NES consensus redefined by structures of PKI-type and Rev-type nuclear export signals bound to CRM1. Nat. Struct. Mol. Biol. 17, 1367–1376 (2010).

40. Lange, A. et al. Classical nuclear localization signals: definition, function, and interaction with importin α. J. Biol. Chem. 282, 5101–5105 (2007).

41. Abramson, J. et al. Accurate structure prediction of biomolecular interactions with AlphaFold 3. Nature 630, 493–500 (2024).

42. Evans, R. et al. Protein complex prediction with AlphaFold-Multimer. bioRxiv 2021.10.04.463034 (2022) doi:10.1101/2021.10.04.463034.

43. Alberti, S., Gladfelter, A. & Mittag, T. Considerations and Challenges in Studying Liquid-Liquid Phase Separation and Biomolecular Condensates. Cell 176, 419–434 (2019).

44. Riback, J. A. et al. Composition-dependent thermodynamics of intracellular phase separation. Nature 581, 209–214 (2020).

45. Wang, J. et al. A Molecular Grammar Governing the Driving Forces for Phase Separation of Prion-like RNA Binding Proteins. Cell 174, 688–699.e16 (2018).

46. von Bülow, S., Tesei, G., Zaidi, F. K., Mittag, T. & Lindorff-Larsen, K. Prediction of phase-separation propensities of disordered proteins from sequence. Proc. Natl. Acad. Sci. 122, e2417920122 (2025).

47. Ibáñez de Opakua, A., Pantoja, C. F., Cima-Omori, M.-S., Dienemann, C. & Zweckstetter, M. Impact of distinct FG nucleoporin repeats on Nup98 self-association. Nat. Commun. 15, 3797 (2024).

48. Nott, T. J. et al. Phase Transition of a Disordered Nuage Protein Generates Environmentally Responsive Membraneless Organelles. Mol. Cell 57, 936–947 (2015).

49. Brady, J. P., et al. Structural and hydrodynamic properties of an intrinsically disordered region of a germ cell-specific protein on phase separation. Proc. Natl. Acad. Sci. 114, E8194–E8203 (2017).

50. Schuster, B. S. et al. Identifying sequence perturbations to an intrinsically disordered protein that determine its phase-separation behavior. Proc. Natl. Acad. Sci. 117, 11421–11431 (2020).

51. Martin, E. W. et al. Valence and patterning of aromatic residues determine the phase behavior of prion-like domains. Science 367, 694–699 (2020).

52. Choi, J.-M., Holehouse, A. S. & Pappu, R. V. Physical Principles Underlying the Complex Biology of Intracellular Phase Transitions. Annual Review of Biophysics vol. 49 107–133 (2020).

53. Holehouse, A. S. & Alberti, S. Molecular determinants of condensate composition. Mol. Cell 85, 290–308 (2025).

54. Bremer, A. et al. Deciphering how naturally occurring sequence features impact the phase behaviours of disordered prion-like domains. Nat. Chem. 14, 196–207 (2022).

55. Banerjee, P. R., Milin, A. N., Moosa, M. M., Onuchic, P. L. & Deniz, A. A. Reentrant Phase Transition Drives Dynamic Substructure Formation in Ribonucleoprotein Droplets. Angew. Chem. Int. Ed. 56, 11354–11359 (2017).

56. Krainer, G. et al. Reentrant liquid condensate phase of proteins is stabilized by hydrophobic and non-ionic interactions. Nat. Commun. 12, 1085 (2021).

57. Ho, J., Jain, A. & Abbeel, P. Denoising Diffusion Probabilistic Models. in Advances in Neural Information Processing Systems (eds Larochelle, H., Ranzato, M., Hadsell, R., Balcan, M. F. & Lin, H.) vol. 33 6840–6851 (Curran Associates, Inc., 2020).

58. Song, Y., et al. Score-Based Generative Modeling through Stochastic Differential Equations. in International Conference on Learning Representations (2021).

59. Esser, P. et al. Scaling rectified flow transformers for high-resolution image synthesis. in Proceedings of the 41st International Conference on Machine Learning (JMLR.org, 2024).

60. Loshchilov, I. & Hutter, F. Decoupled Weight Decay Regularization. in International Conference on Learning Representations (2019).

61. Lin, Z. et al. Evolutionary-scale prediction of atomic-level protein structure with a language model. Science 379, 1123–1130 (2023).

62. Gretton, A., Borgwardt, K. M., Rasch, M. J., Schölkopf, B. & Smola, A. A Kernel Two-Sample Test. J. Mach. Learn. Res. 13, 723–773 (2012).

63. Stringer, C. & Pachitariu, M. Cellpose3: one-click image restoration for improved cellular segmentation. Nat. Methods 22, 592–599 (2025).

